# In Silico Transcriptome-based Screens Identify Epidermal Growth Factor Receptor Inhibitors as Therapeutics for Noise-induced Hearing Loss

**DOI:** 10.1101/2023.06.07.544128

**Authors:** Sarath Vijayakumar, Joe A. DiGuiseppi, Jila Dabestani, William G. Ryan, Rene Vielman Quevedo, Yuju Li, Jack Diers, Shu Tu, Jonathan Fleegel, Cassidy Nguyen, Lauren M. Rhoda, Ali Sajid Imami, Ali Abdul-Rizaq Hamoud, Sándor Lovas, Robert McCullumsmith, Marisa Zallocchi, Jian Zuo

## Abstract

Noise-Induced Hearing Loss (NIHL) represents a widespread disease for which no therapeutics have been approved by the Food and Drug Administration (FDA). Addressing the conspicuous void of efficacious in vitro or animal models for high throughput pharmacological screening, we utilized an in silico transcriptome-oriented drug screening strategy, unveiling 22 biological pathways and 64 promising small molecule candidates for NIHL protection. Afatinib and zorifertinib, both inhibitors of the Epidermal Growth Factor Receptor (EGFR), were validated for their protective efficacy against NIHL in experimental zebrafish and murine models. This protective effect was further confirmed with EGFR conditional knockout mice and EGF knockdown zebrafish, both demonstrating protection against NIHL. Molecular analysis using Western blot and kinome signaling arrays on adult mouse cochlear lysates unveiled the intricate involvement of several signaling pathways, with particular emphasis on EGFR and its downstream pathways being modulated by noise exposure and Zorifertinib treatment. Administered orally, Zorifertinib was successfully detected in the perilymph fluid of the inner ear in mice with favorable pharmacokinetic attributes. Zorifertinib, in conjunction with AZD5438 – a potent inhibitor of cyclin dependent kinase 2 – produced synergistic protection against NIHL in the zebrafish model. Collectively, our findings underscore the potential application of in silico transcriptome-based drug screening for diseases bereft of efficient screening models and posit EGFR inhibitors as promising therapeutic agents warranting clinical exploration for combatting NIHL.

**Highlights:**

1. In silico transcriptome-based drug screens identify pathways and drugs against NIHL.
2. EGFR signaling is activated by noise but reduced by zorifertinib in mouse cochleae.
3. Afatinib, zorifertinib and EGFR knockout protect against NIHL in mice and zebrafish.
4. Orally delivered zorifertinib has inner ear PK and synergizes with a CDK2 inhibitor.

## INTRODUCTION

Noise-induced hearing loss (NIHL) is one of the most common forms of sensorineural hearing loss and occupational hazards in modern society worldwide (*1–6*). The impact of hearing loss on our society is such that one in six Americans may exhibit some degree of hearing loss (*7, 8*). Occupational noise exposure has been attributed to about 16% of disabling hearing loss worldwide (*9*). Approximately 1 billion young adults and adolescents are at risk for NIHL due to recreational exposure to noise via personal audio systems, loud music in clubs, and at music concerts (World Health Organization, 2015). NIHL significantly affects the military and veterans. Centers for Disease Control and Prevention reported that military veterans have four times higher risk of developing severe hearing loss compared to age- and occupation-matched civilians (*10*). NIHL has a negative impact on the quality of life and carries a significant financial burden on affected individuals (*11*). The total economic cost of hearing loss, including NIHL, is over $750 billion annually worldwide (*12*). Hearing loss has been implicated as a potential risk factor for accelerated cognitive decline and impairment in the increasingly socially isolated elderly (*13–15*). It has also been linked to the development of depression in some individuals (*13*). Despite the enormous societal impact, there are no drugs that are approved by the FDA to prevent or recover NIHL. Currently, hearing aids that amplify the sound and cochlear implants are the mainstay approaches to treating hearing loss. Depending on the intensity and duration of the noise exposure, acoustic trauma can cause damage to the cochlear hair cells, supporting cells, hair cell synapses, or spiral ganglion neurons, all of which can lead to hearing impairment (*16, 17*). Significant advances have been made in our understanding of the cellular and molecular processes involved in the pathophysiology of NIHL, including but not limited to glutamate excitotoxicity, oxidative stress, imbalance of ions in the endolymph, inflammation, and microcirculation changes in the stria vascularis (*18*). Since NIHL is a predictable form of hearing loss, it is feasible to prevent it by inhibiting cochlear cell death or promoting cochlear cell survival. Despite extensive research in recent years, most candidate compounds currently in pre-clinical and clinical trials are related to antioxidants, vitamins, and glutathione metabolism, and their effectiveness remains unclear (*19, 20*).

Over the last two decades, high-throughput screening (HTS) has become a standard approach in drug discovery. Recently, HTS has uncovered small molecule otoprotectant candidates (*21–27*). These chemical phenotypic screenings are unbiased in that they explore diverse biological pathways that prevent cisplatin- or antibiotic-induced cochlear cell death in cell lines, explants, or zebrafish models. Unfortunately, such drug screens for noise protection cannot be easily applied to cell lines, explants, or zebrafish models, since these assays cannot accurately simulate the inner ear milieu during noise exposure for HTS. Thus, we used a computational drug discovery approach to identify the biological pathways and drugs of most interest to the prevention and treatment of NIHL. Drug development strategies based on transcriptomics are advantageous in that they do not require a large amount of a priori knowledge pertaining to particular diseases or drugs (*28, 29*). In silico screening using the connectivity map (CMap) requires a gene expression profiling database of small molecules to be compared with the gene expression signatures of a disease or condition such as NIHL (*30*). The library of integrated network-based cellular signatures (LINCS) L1000 dataset currently has over a million gene expression profiles in small molecule treated cell lines (*31*). Drug candidates can be predicted by comparing the LINCS L1000 CDS^2^ perturbation signatures and the disease specific signatures extracted from the gene expression omnibus (*32*). By comparing published mouse cochlear gene expression profiles in NIHL and the LINCS L1000 dataset, we identified 22 candidate pathways and 64 candidate drugs protective against NIHL. Interestingly, among the top hits are tyrosine kinase inhibitors that target epidermal growth factor receptor (EGFR) and human epidermal growth factor receptor 2 (HER2). EGFR is a member of the epidermal growth factor receptor family that includes HER1 (erbB1, EGFR), HER2 (erbB2, NEU), HER3 (erbB3), and HER4 (erbB4), which activate and regulate diverse processes including cell survival, proliferation, differentiation, and migration (*33*). EGFR is widely expressed on the surface of mammalian epithelial, mesenchymal, and neuronal cells. EGFR transcripts have been detected in postnatal rat cochlear organotypic cultures, in multiple cell types in neonatal mouse cochleae, including both inner and outer hair cells, spiral ganglion neurons, and Deiters’ and Hensen cells (*34–36*). ErbB2, ErbB3, and ErbB4 immunolabeling is present in the cochlear and vestibular sensory epithelia of adult chinchilla (*37*). While studies have looked at the role of ErbB2 in the survival and regeneration of hair cells in the cochlea (*38, 39*), inhibition of EGFR as a therapeutic intervention for noise-induced hearing loss has not been investigated. In this study, we focused on providing “proof of concept” for the future drug development of EGFR inhibitors as otoprotective drugs. Our studies provide a robust and promising example of using in silico transcriptome-based screens for therapeutics to treat a common disease that is difficult to perform drug screens.

## RESULTS

### Generation of transcriptome-based cochlear signatures for acoustic trauma

We used three sets of published in vivo transcriptome data (*40–42*) where differentially expressed genes (DEGs) in adult mouse cochleae with or without noise-exposure were obtained using microarray, NextGen RNA-seq and single cell RNA-seq analyses (**Fig. 1A**; see Methods). For the Gratton et al. and Maeda et al. datasets, DEGs were first identified using GEO2R and ranked based on p-value and logFC: DEGs with a |logFC| > 2 and p-value < 0.05 were considered DEGs of interest to be included in further in silico analyses. For the Milon et al. dataset, DEGs for outer hair cells, supporting cells, spiral ganglion neurons, and stria vascularis from the published lists were utilized without any further processing. In this dataset, a gene with absolute fold change > 1.2 and a false discovery rate (FDR) q-value <0.05 was considered differentially expressed. Pathway analysis was performed on all DEGs of interest using ShinyGO Enrichment analysis, and Gene Ontology (GO) enrichment pathways were ranked based on enrichment FDR values. LINCS L1000CDS2 was then compared for each GO enrichment pathway with at least three upregulated and three downregulated DEGs of interest, which resulted in a list of drug perturbations that could mimic or reverse the input gene expression in cancer cell lines. Drug perturbations were ranked based on overlapping scores with the input gene list. Drugs with the highest overlapping scores were identified from each pathway, and drugs that targeted multiple significant pathways were considered more promising and advanced for in vivo studies. In total, 246 novel drug perturbations were identified for the prevention and/or treatment of NIHL based on microarray and RNA-seq transcriptomic analysis.

**Fig. 1.**
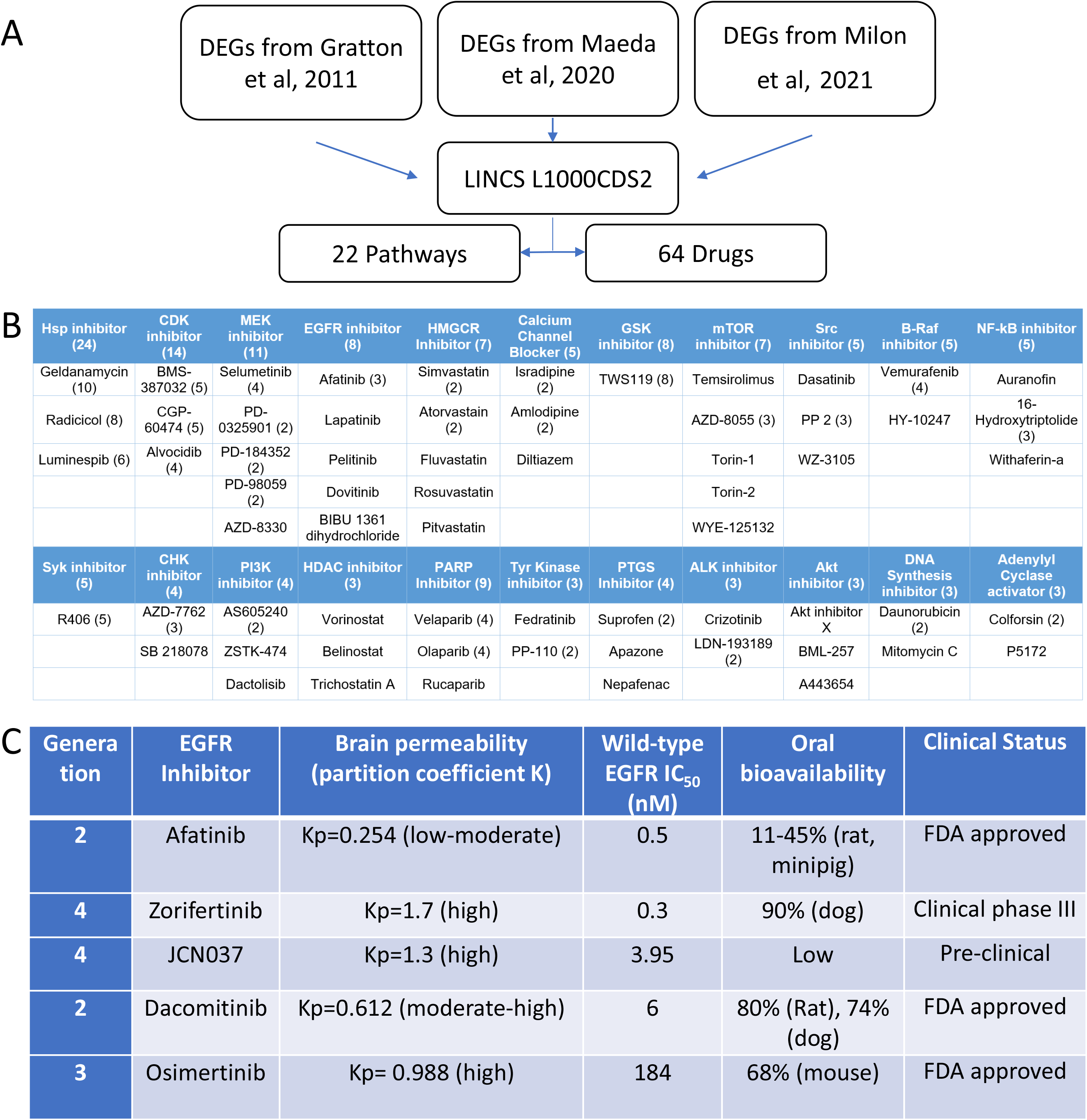
(**A**) Workflow on in silico screens for pathways and drugs to protect against NIHL. (**B**) Top biological pathways and drug candidates identified against NIHL. (**C)** Top five EGFR inhibitors with proper safety and PK/PD profiles.

### Ranking biological pathways and candidate drugs by their otoprotectant potential

After compiling the list of 246 novel drug perturbations, we determined each drug’s targets, predicted mechanisms of action, number of hits, and phase of FDA approval. L1000 Fireworks Display (L1000FWD) was used in conjunction with literature queries to determine which biological pathways are affected by each drug. Once the literature review was completed and the effected biological pathway was known for each drug, the pathways were ranked based on the number of hits found in each pathway; pathways with the most hits are more strongly related to NIHL protection than pathways with fewer hits. This method of pathway ranking assumes that all the pathways involved will reverse the damage caused by noise rather than reversing possible protective pathways. Pathways with at least three hits were considered pathways of interest which require further study. Using this threshold, 22 biological pathways and 64 drugs targeting those pathways were identified (**Fig. 1B**).

Once the significant pathways related to NIHL were identified, our next step was to determine the drugs to be advanced for testing in animal models against noise exposure. We first focused on FDA-approved small molecule inhibitors in high-ranking pathways of MEK, EGFR, mTOR, Src, and HDAC, with excellent reported therapeutic properties and publications on other indications (i.e., PK/PD, maximum tolerated dose [MTD], blood-brain-barrier [BBB] permeability (*43*), preclinical and clinical status). Among these inhibitors and pathways, many are known to be involved in NIHL (i.e., Braf, MEK, HDAC, and CDK) (*22, 27, 44, 45*), further validating our in silico screening strategies.

We selected the epidermal growth factor receptor (EGFR) inhibitors among our top otoprotectant candidates against NIHL from our noise transcriptomic LINCS analyses. Because EGFR is upstream of multiple pathways involved in NIHL, we reasoned that EGFR inhibitors may have better effects than those with individual pathways (i.e., Braf, MEK, HDAC, and CDK). We further searched additional EGFR inhibitors in the literature for high affinity to wild-type EGFR, high partition coefficient (Kp) for brain permeability, high bioavailability and already in clinical trials (**Fig. 1C**). Among the top five EGFR inhibitors identified, afatinib, a 2^nd^ generation EGFR inhibitor approved by FDA for cancer treatment, was the top-ranking drug in the enrichment analyses that was included in multiple pathways involved in the pathogenesis of NIHL. Zorifertinib (AZD3759) is a 4^th^ generation, BBB penetrating EGFR inhibitor currently in cancer clinical trials. Molecular docking analyses further support that afatinib, zorifertinib and other top EGFR inhibitors target the active kinase sites of EGFR (**Supplemental Fig. 1**).

### EGFR signaling in the adult mammalian cochlea

To provide evidence of expression of EGFR and downstream signaling pathways in the adult cochlear supporting cells and hair cells, we analyzed our published single cell (sc)RNA-seq datasets of P28 and P33 mouse cochlear supporting cells and hair cells (**Fig. 2; Supplemental Fig. 2**) (*46, 47*). Representative EGF ligands (Egf, Tgfa, Hbegf, Nrg1, Nrg4, and Btc) are all expressed in P28 and P33 supporting cells; EGFR family members (EGFR, ErbB2, ErbB3, and ErbB4) are expressed in P28 and P33 supporting cells and hair cells; and EGFR downstream signaling pathway genes (Akt/Pi3k, Erk/Mapk, Stat3 and Plc) are expressed in P28 and P33 supporting cells and hair cells. These results are consistent with immunostaining results in adult cochleae (*48, 49*).

**Fig. 2.**
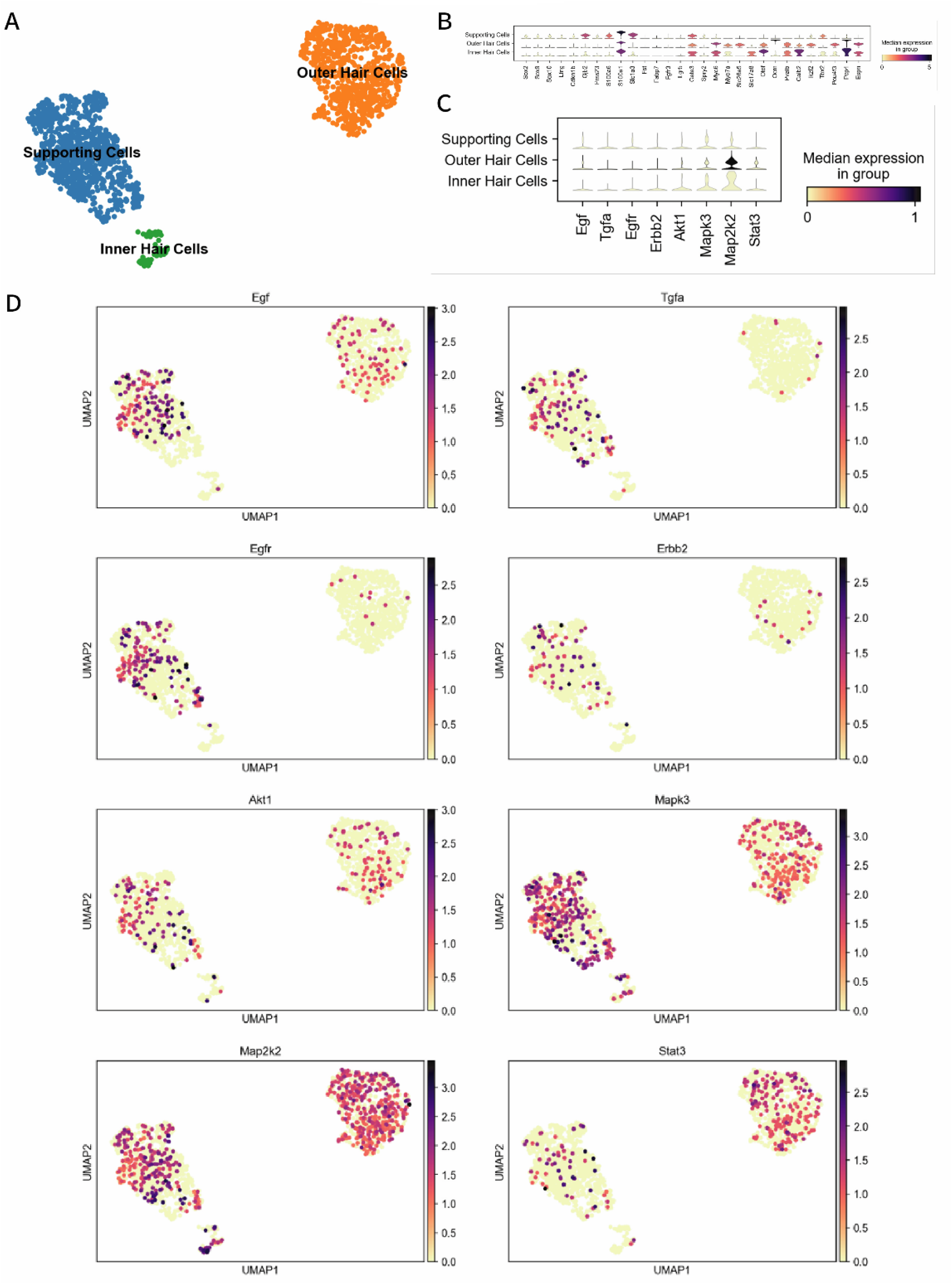
EGFR signaling in the adult mammalian cochlea. Single-cell RNA sequencing reveals EGFR ligands, receptors, and nstream targets in hair and supporting cells. (**A**) UMAP plot with Leiden clustering showing supporting cell (blue), outer hair cell nge), and inner hair cell (green) clusters in adult P28 C57BL6 mice. (**B**) Stacked violin plot showing hair and supporting cell kers in the clusters. (**C**-**D**) Expression levels and distribution of ligands (Egf and Tgfa), receptors (Egfr and Erbb2), and nstream targets (Akt1, Erk1/Mapk3, Map2k2, and Stat3) in the hair and supporting cell clusters. All single cell RNA sequencing ysis done using Scanpy 1.9.3 (*110*). The code used: https://nbviewer.org/github/renevq/jupyter-books/blob/main/Hang_data.ipynb. Similar results were obtained based on our published dataset (*46*).

### EGFR inhibitors protect against hair cell excitotoxicity in zebrafish

Because there is no established mammalian cochlear explant assay that mimics noise injury, we adopted a zebrafish model that mimics hair cell damage due to excitotoxicity (*50*). Previous studies found that NIHL may be caused, in part, by glutamate excitotoxicity (*51–53*). Ionotropic glutamate receptor agonists have been used to mimic noise exposure in zebrafish larvae (*50*). We therefore tested the efficacy of top five EGFR inhibitors (afatinib, zorifertinib, osimertinib, JCN037, and dacomitinib) to protect against kainic acid (KA)-induced excitotoxicity in this zebrafish lateral line neuromast model. This zebrafish excitotoxicity assay is by no means ideal for drug validation for NIHL but can serve as a high-throughput model for drug screen and validation prior to testing in mammalian models. Importantly, if a drug is protective in both zebrafish and mouse models of NIHL, it will provide evidence of conservation of mechanisms of action of the drug across species supporting human clinical use.

Five-to-six days post fertilization (dpf) *Tg (brn3c:GFP)* zebrafish larvae were used for the assay. Fish were incubated with 500 mM of KA for one hour followed by various concentrations of the different EGFR inhibitors (**Fig. 3A-E**). Except for JCN037 (**Fig. 3B**), all the EGFR inhibitors were protective at more than one concentration. To provide evidence that EGFR inhibitors act through the EGFR signaling pathway, we knocked down (KD) the EGF ligand (EGFL) in zebrafish (**Fig. 3J**) and tested protection against N-methyl-D-aspartate (NMDA) (**Fig. 3F**) and KA (**Fig 3G- I**). As expected, the incubation of the EGFL knockdown animals with the EFGR inhibitors did not confer protection against excitotoxic damage (**Fig. 3F-I**), suggesting that the protective effect is mediated via EGFR pathway and that EGFR is the major pathway by which afatinib and zorifertinib protect against excitotoxicity. The use of two different EGFR inhibitors in combination with EGFL KD strongly points to the involvement of EGFR signaling cascade during hair cell excitotoxicity. To be noted, there were no differences in the number of neuromas hair cells between non-injected, scrambled-injected and EGFL-injected animals, confirming the lack of morpholino off-target effect (**Fig. 3K**).

**Fig. 3.**
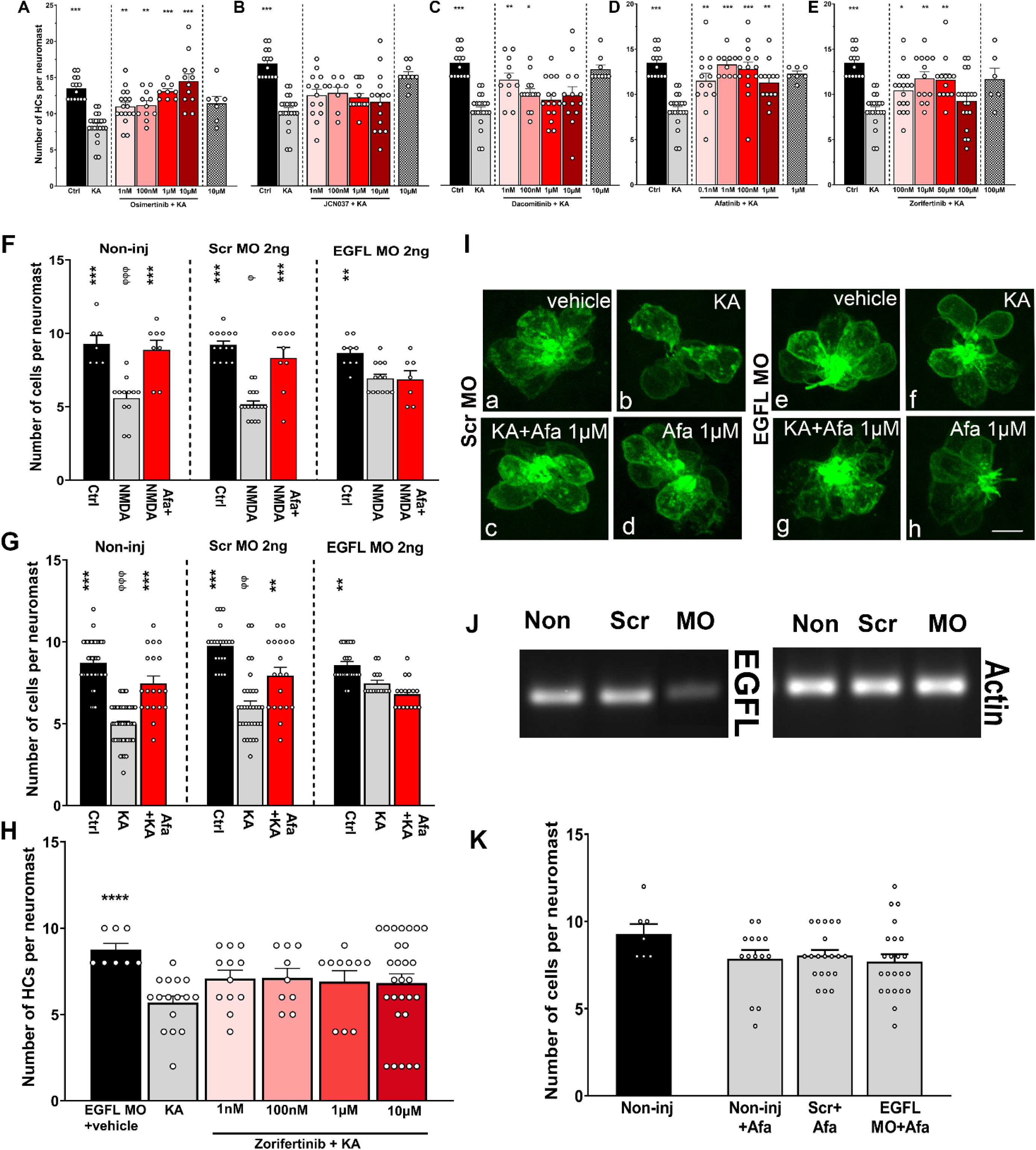
EGFR inhibitors protect against hair cell excitotoxicity in zebrafish. **A-E**: to-six-days post-fertilization zebrafish were incubated with 500 mM of KA for 1 followed by two-hour incubation with different concentrations of EGFR inhibitors. simertinib, **B**: JCN037, **C**: Dacomitinib, **D**: Afatinib, **E**: Zorifertinib. The number of cell is expressed as mean +/- SEM. *P<0.05, **P<0.01, ***P<0.001 versus KA-One-way ANOVA. **F-K:**Zebrafish non-injected or injected with 2ng of mbled or EGF ligand (EGFL) specific morpholinos (Gene Tools). Animals (3dpf) pre-incubated with 500 mM of NMDA (**F**), or KA (**G**) for 1 hour followed by 2 incubation with vehicle or afatinib (1µM). NMDA and KA incubations were used mic noise exposure (40). Quantification of the HCs was performed in SO3, O1 O2 neuromasts. **P<0.01, ***P<0.001 *versus* ototoxin alone. ɸ P<0.05, ɸɸ 01, ɸɸɸ P<0.001 versus the corresponding control (One-way ANOVA). **H:** afish EGFL KDs incubated with KA (500 mM) followed by different concentrations orifertinib. ***P<0.01 versus KA alone (One-way ANOVA). **I:** Representative es of scrambled and EGFL morphants with the different treatments. GFP in denotes the neuromast hair cells. Ctrl: control, KA: Kainic acid incubation, fa 1mM: Kainic acid and afatinib incubations, Afa 1mM: afatinib-only incubation. le treated EGFL morphants did not show any significant differences compared to njected or scrambled controls. **J:** Confirmation of EGFL knockdown by RT-PCR rambled and EGFL morphants. **K:** Hair cell quantification in the different hants under baseline conditions (no excitotoxicity). Hair cell quantification is ssed as mean ± SEM.

### Afatinib and zorifertinib protect against noise-induced hearing loss in mice

Otoprotective effects of afatinib and zorifertinib were further evaluated in a mouse model of acute NIHL. We used a previously established permanent threshold shift (PTS) FVB mouse model (*27*) to test the otoprotective effect. After baseline auditory function evaluation using auditory brainstem response (ABR) and distortion product otoacoustic emission (DPOAE) measurements, mice were exposed to 100 dB SPL 8-16 kHz octave band noise for 2 hours. Experimental animals were given either afatinib (20 mg/kg IP delivery) or zorifertinib (15 mg/kg oral delivery) one day before the noise exposure and then continued for following 4 days. Treatment with afatinib or zorifertinib significantly attenuated permanent threshold shifts (PTS) 14 days post-noise exposure compared to the vehicle treated group (**Fig. 4A-D**). Drug only treated animals did not show any significant difference in the threshold shifts. ABR threshold shifts were significantly attenuated at 16, 22.6, 32, 45.2 and 64 kHz frequencies for both drug treatment groups (**Fig. 4A & C**). DPOAE threshold shifts were most prominently protected in afatinib treated animals at 22.6 kHz (**Fig. 4B**). Although zorifertinib treated animals also showed lower DPOAE threshold shifts compared to the vehicle treated group, it did not reach statistical significance (**Fig. 4D**). In addition, significant differences were observed at multiple stimulus levels in ABR wave 1 amplitude input-output function at both 16 and 22.6 kHz frequencies in the afatinib treated group (**Fig. 4E & F**).

**Fig. 4.**
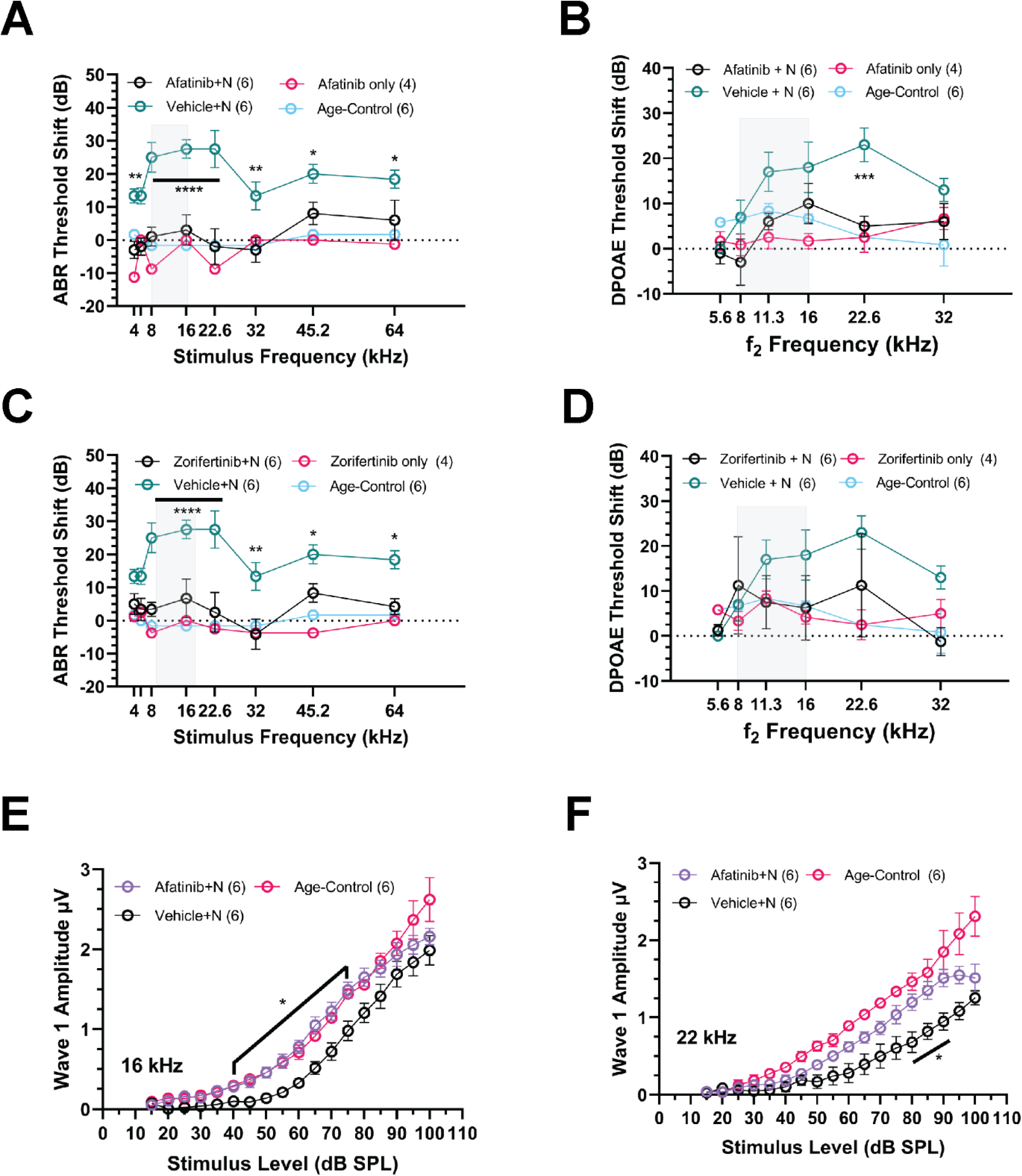
Afatinib and zorifertinib protect against noise-induced hearing loss mice. Adult FVB mice (4-7 weeks old) were exposed to noise trauma (8-16kHz se band at 100 dB SPL for 2hrs) indicated by the shaded box in the figures. ABR threshold shifts 2 weeks after the noise-exposure. Animals treated with 4 es of afatinib 20 mg/kg/day IP (magenta) showed significant difference in the eshold shifts at all tested frequencies when compared to those treated with the icle (teal). **(B)** DPOAE thresholds (a function of the outer hair cells of the hlea; defined as the lowest level of f2 that produced an emission amplitude of dB SPL and was also 6 dB higher than the corresponding noise floor). nificant difference in threshold were observed at 22.6kHz (p=0.0005). **(C)** ABR eshold shifts 2 weeks after noise-exposure. Animals treated with 5 doses of ifertinib 15mg/kg oral gavage showed significant protection at all tested quencies above 16kHz (8,16 and 22 kHz p<0.0001 ;32kHz p=0.0011, 45.2kHz 0.0447 and 64kHz p=0.0112). **(D)** DPOAE threshold difference did not reach tistical significance. (**E & F**) ABR Wave-1 amplitudes for 16kHz and 22 kHz mulus respectively showed significantly larger amplitudes at suprathreshold els in the afatinib treated animals compared to the vehicle treated (p=0.0085 – 498 for afatinib and p=0.0233 – 0.0352 at 80-90 dB SPL for zorifertinib). Two y ANOVA, Holm-Šidák post hoc test. Data are presented as mean ± SEM, 4-6/group.

### Afatinib and zorifertinib protect against cochlear synaptopathy in mice

In the acute NIHL mouse model, moderately loud noise exposure doesn’t cause hair cell loss but is associated with permanent loss of cochlear afferent synapses on the IHCs (*27, 54, 55*). Therefore, we examined synaptic ribbons in the IHCs in the 16-22 kHz region of the cochlea (**Supplemental Fig. 3**). Without noise, we found no difference in the number of CtBP2 presynaptic puncta among the zorifertinib, afatinib, and age-matched control groups. However, after noise exposure, there were significantly more CtBP2 presynaptic puncta in the zorifertinib and afatinib treated groups compared to the control, suggesting zorifertinib and afatinib protect against noise induced cochlear synaptopathy.

### Conditional knockout of EGFR protects against NIHL in mice

To further validate the protective effect of inhibition of EGFR signaling against NIHL, we generated a conditional knockout (cKO) mouse strain of EGFR by crossing EGFR floxed mice with Pax2-Cre mice. The Pax2-Cre; EGFR cKO mice showed normal hearing thresholds and cochlear morphology (data not shown) as their wildtype littermates (**Fig. 5A**).

**Fig. 5.**
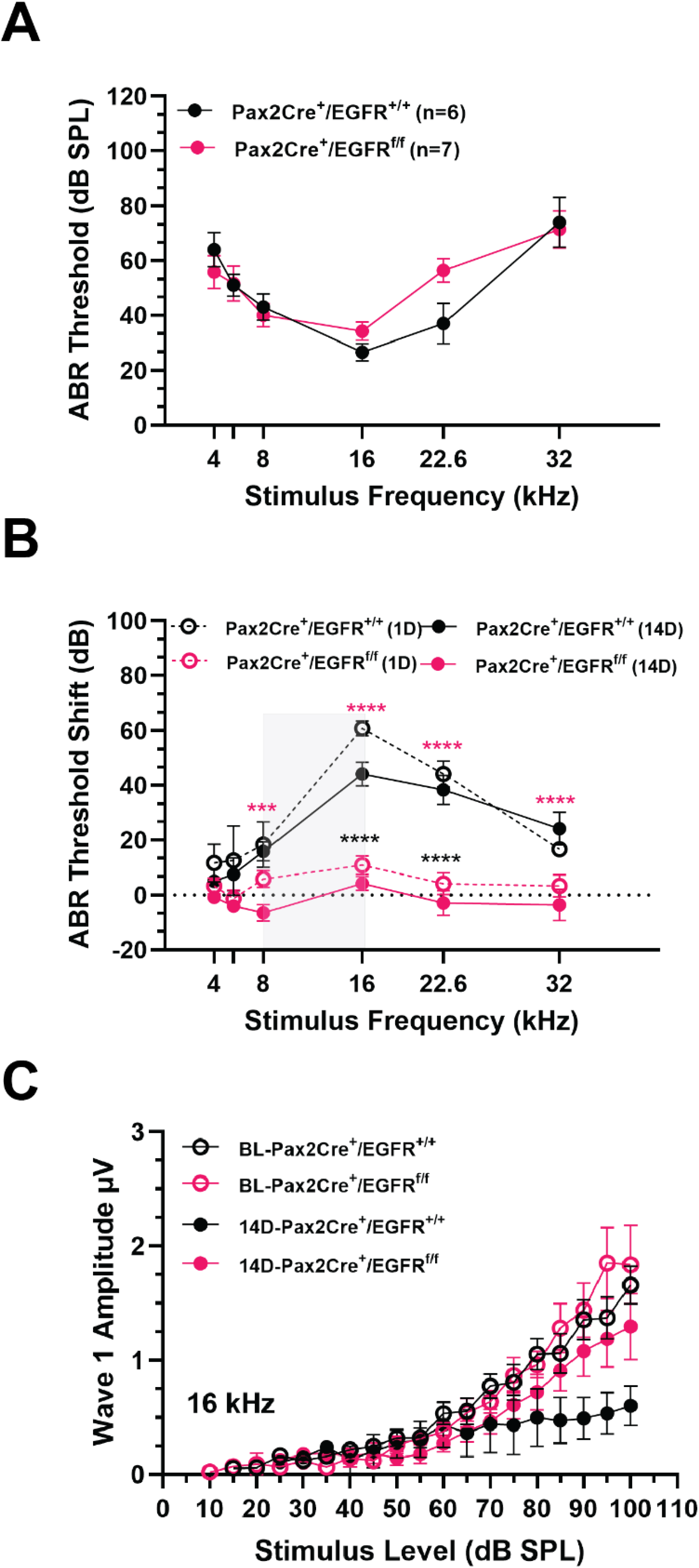
Conditional knockout of EGFR shows protection against NIHL in mice. (**A**) Baseline (pre-noise exposure) ABR thresholds of Pax2Cre; EGFR^flox/flox^ mice and Pax2Cre; EGFR^+/+^ control littermates. No significant differences were detected between cKO and WT. n=6-7. (**B**) Pax2Cre; EGFR^flox/flox^ mice (n=6) exhibited significantly smaller ABR threshold shifts (both 1-day (1D) for TTS and 14-days (14D) for PTS) than wild-type (WT) littermate controls (Pax2Cre; EGFR^+/+^; n=6) C) ABR Wave-1 amplitudes for 16kHz stimulus respectively showed larger amplitudes at suprathreshold levels in the cKO compared to the WT but did not reach statistical significance. Two-way ANOVA, Holm-Šidák post hoc test, ***** p<0.0001, *** p=0.0007, black (1D), magenta (14D). Data are presented as mean ± SEM.

To test whether the Pax2-Cre; EGFR cKO mice were protected against NIHL, we exposed the mice to 2 hrs of 100 dB SPL 8-16 kHz octave band noise, like the noise exposure in the afatinib and zorifertinib treated groups. cKO mice displayed significantly smaller ABR threshold shifts at 8, 16, 22.6 and 32 kHz frequencies on both 1 day and 14 days after the noise exposure compared to the wildtype littermate controls (**Fig. 5B**). Wave 1 amplitudes of ABRs at 16 kHz were larger but did not reach significance in the cKO mice (**Fig. 5C**). These results are similar to the protection against NIHL offered by pharmacological inhibition of EGFR signaling (afatinib and zorifertinib; **Fig. 4**).

### EGFR signaling is activated in the mouse cochlea following noise exposure and attenuated by zorifertinib

To determine whether EGFR signaling is functional in the adult cochlea, we treated wild type FVB mice with zorifertinib (oral gavage at 15 mg/kg/day) a day before and at 1 hr during noise trauma. We exposed the mice to noise trauma (100dB SPL 8-16 kHz for 2 hrs.) and performed immunoblot analysis for downstream signaling targets of EGFR on cochlear lysates collected 30 minutes after the noise trauma. We found that ERK and AKT phosphorylation was induced 30 min after noise exposure, but significantly reduced by the drug treatments (**Fig. 6A- D**). Our results corroborate with previous publications on noise induced activation of ERK and AKT pathways in similar post-exposure time points (*56–59*). Together, these results strongly indicate that EGFR signaling is functional and responsive to noise exposure. Most importantly, EGFR inhibitors are effective in mitigating the noise-induced activation of AKT/ERK signaling pathways in the adult mouse cochlea in vivo.

**Fig. 6.**
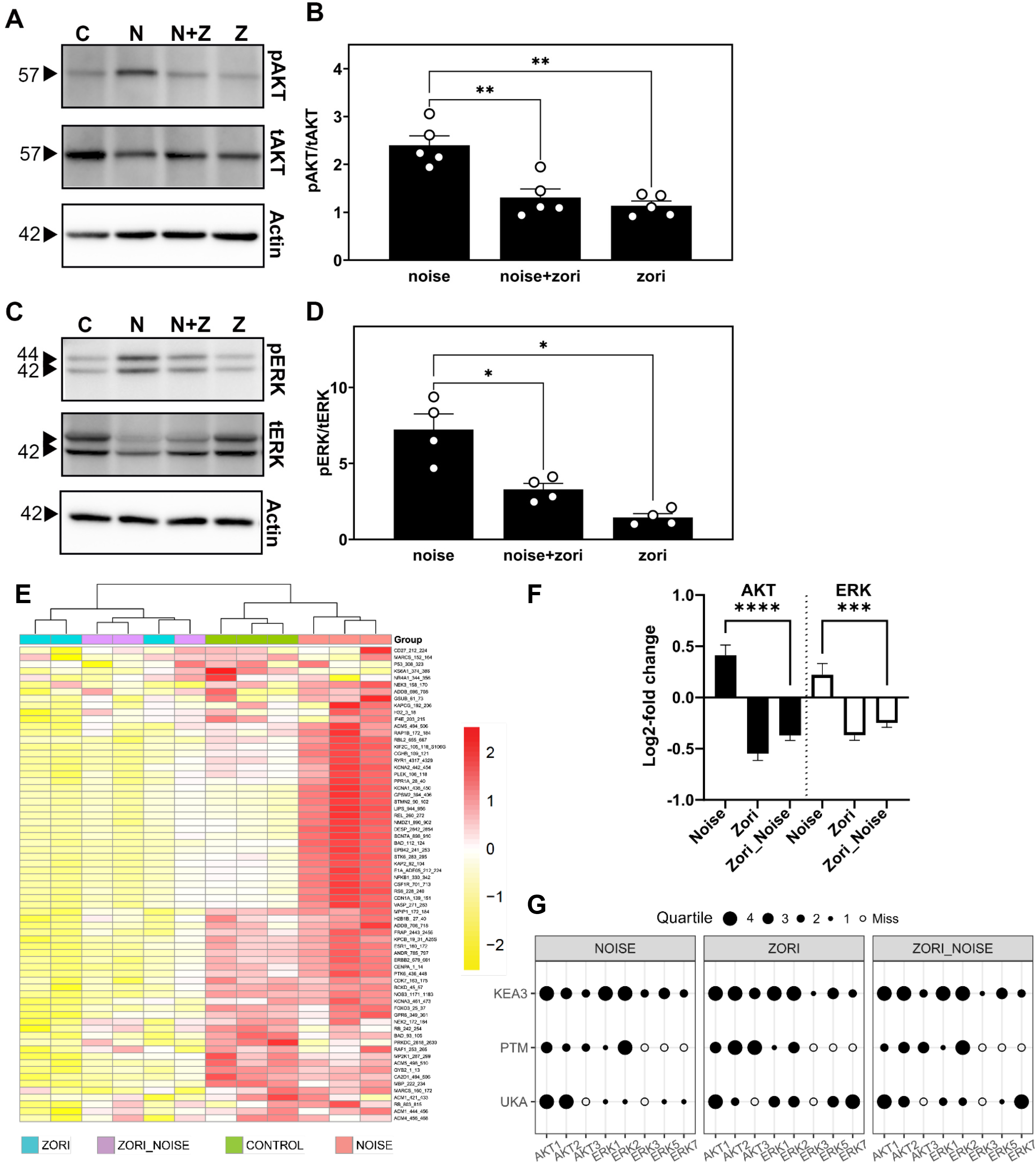
EGFR signaling is activated in the mouse cochlea following noise osure and attenuated by zorifertinib. (**A-D**) Western blotting was performed rgan of Corti lysates from mice 30 min after exposed to noise trauma (8-16kHz e band at 100 dB SPL for 2hrs) and pretreated with either the drug (zorifertinib g/kg oral) or the vehicle (normal saline or methyl cellulose) one day before and during the noise trauma to detect the phosphorylation status (p phosphorylated total) of two downstream effectors of EGFR (AKT and ERK). Zorifertinib eatment significantly decreased the noise induced increase of AKT and ERK phorylation. (A and C) The sizes of bands are labeled in kD. C – untreated and xposed control; V+N –vehicle + Noise; Z+N – zorifertinib + Noise; Z – ertinib only. Actin: a loading control. (B and D) Ratios of phosphorylated vs total eins normalized to actin loading controls. Data are normalized to eated/exposed control mice as a ratio of 1 and presented as mean ± SD. N = 4-5 gical replicates and each dot represents one mouse in each group. ANOVA, y’s post hoc. * p≤0.05, ** p≤0.01. (**E-G**) Zorifertinib suppresses noise-activated R signaling pathways in mouse cochleae. (**E**) Heatmap of phosphorylation ity at the reporter peptides on the kinome panel (STK Pamchip; n=3 chips run in nical triplicate). Cochlear sensory epithelial lysates were collected 30-min post e exposure from control, noise-exposed, zorifertinib-treated (Zori), and ertinib-treated plus noise-exposed (Zori_Noise) adult FVB mice. Each row esents a peptide. The protein kinase activity measured as phosphorylation of ted peptides on the chip is scaled with red being higher activity and yellow ating lower activity. (**F**) Log2-fold activity changes of reporter peptides “mapped” utative targets of AKT and/or ERK kinase families (24 and 25 peptides, ectively) expressed as mean +/- standard deviation. Average of triplicates was for each peptide per condition. N/C: Noise vs Control; Z/C: Zorifertinib vs rol; N+Z/C: Noise + Zorifertinib vs Control. Bar: SEM; ****: P<0.0001, ***: 001 Two-way ANOVA. (**G**) Identification of specific protein kinases using plementary software packages (details in supplement). Selected AKT and ERK y members were identified as candidate “hits” using deconvolution strategies specify protein kinases that are contributing to the phosphorylation signal seen ss all reporter peptides on the STK chip. Upstream Kinase Analysis (UKA), translational Modification Signature Enrichment Analysis (PTM-SEA), and se Enrichment Analysis 3 (KEA3) were used to identify specific kinases within AKT and ERK families using online phosphosite mapping databases. Data were alized and grouped as quartiles (1-4 black dots) for visualization using our oke Creedenzymatic R package.

To further elucidate the mechanisms of action by zorifertinib against noise trauma in the adult mouse cochlea, we used a novel kinome array to compare kinase signaling pathways in the cochlear lysates in mice treated with zorifertinib and/or noise trauma (see Methods). Specifically, a serine/threonine kinase (STK) Pamchip containing 144 immobilized peptides which were used as a readout of kinase activity (**Supplemental Fig. 4**). Adult FVB mice were treated with zorifertinib (oral gavage at 15 mg/kg/day one day prior and 1 hr during noise trauma), exposed to noise (100 dB SPL 8-16 kHz for 2 hrs), and cochlear lysates collected 30 min post-noise exposure, in conditions identical to Western blot analysis. We found that technical triplicates of each condition were highly reproducible (**Fig. 6E; Supplemental Figs. 5-14**), zorifertinib suppressed multiple signaling pathways, and signaling downstream of EFGR was higher in noise exposed conditions but treatment with zorifertinib suppressed noise-induced activation of AKT/ERK signaling activities 30-min post-noise exposure (**Fig. 6F**). Using various deconvolution strategies (Upstream Kinase Analysis [UKA], Post-translational Modification Signature Enrichment Analysis [PTM-SEA], and Kinase Enrichment Analysis 3 [KEA3]; see Methods), we identified selected AKT and ERK family members as candidate “hits” that are contributing to the phosphorylation signal seen across all reporter peptides on the STK chip (**Fig. 6G**). These results are consistent with our Western results (**Fig. 6A-D**) and previous publications under similar conditions, thus confirming that inhibition of the EGFR signaling pathway is a mechanism of action by zorifertinib against NIHL in the adult mouse cochlea.

### Pharmacokinetics of orally delivered zorifertinib in mouse inner ear perilymph fluids

To validate orally delivered zorifertinib crosses the blood labyrinth barrier (BLB) of the inner ear, we performed perilymph collection of adult FVB mice at various time points after oral gavage of zorifertinib (15 mg/kg). To accurately measure the perilymph concentration of zorifertinib, we performed LC-MS/MS measurement of zorifertinib in ∼1uL perilymph fluid collected from each mouse using crizotinib as an internal standard (IS); the calibration curve showed linearity within the measured range, with a lower limit of quantification (LLOQ) of 5 ng/ml (**Supplemental Fig. 15**). The time course of zorifertinib concentrations in perilymph is shown in **Fig. 7**. The C_max_ was 100 ng/ml and t_max_ was 30 min after oral gavage, while t1/2 was ∼130 min and zorifertinib was cleared from perilymph in 6 hrs. These PK results in the inner ear fluids are consistent with protection of zorifertinib we observed in mice with noise exposure (**Fig. 4C-D**).

**Fig. 7.**
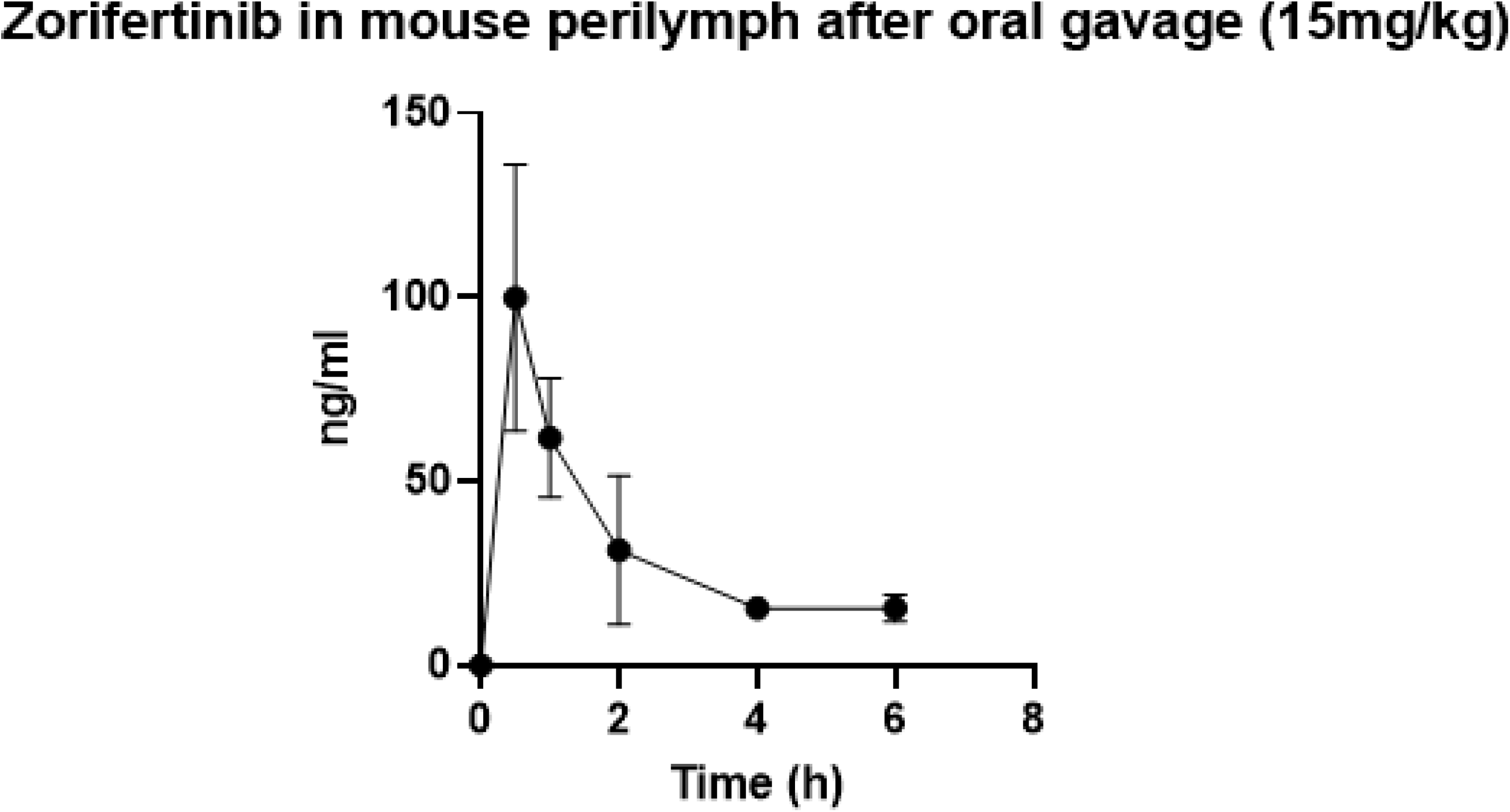
Pharmacokinetics of orally delivered zorifertinib in mouse inner ear perilymph fluids. Measured concentrations of zorifertinib in mouse perilymph 0-6 hours after oral gavage (15 mg/kg; n=3/time point). Error bars indicate SEM. PK properties are calculated: T_max_ = 0.5h (Time of maximum perilymph concentration), T_1/2_ = 2.1h (Half-life), C_max_ = 99.9 +/- 62.6 ng/ml (Maximum concentration), AUC_0-t_ = 189.9 +/- 48.3 ng/ml*h (Area under curve), V_Z_/F = 0.19 (Volume of distribution), Cl/F = 0.06 (mg/kg)/(ng/ml)/h (Clearance).

### Zorifertinib shows synergistic effects with CDK2 inhibitor AZD5438 in zebrafish

Given the multiple pathways involved in pathophysiology of NIHL, we hypothesized that the EGFR inhibitor could synergize with an inhibitor of the CDK2 pathway and thus increase the levels of protection against NIHL. For this purpose, we used the zebrafish model for glutamate excitotoxicity to test the protective effect of zorifertinib in the presence of AZD5438, a CDK2 inhibitor that we have characterized before as an otoprotective compound against NIHL (*22*). Zebrafish larvae were pre-incubated with 500 µM kainic acid or control fish water for 1 hr followed by incubation with a combination of varying doses of zorifertinib and AZD5438. Dose combinations of zorifertinib (1nM) + AZD5438 (50nM), zorifertinib (50nM) + AZD5438 (1nM) and zorifertinib (50nM) + AZD5438 (50nM) showed synergistic protection compared to treatment with the individual drugs (**Fig. 8**). These results demonstrate that zorifertinib and the CDK2 inhibitor, AZD5438, act in synergy against NIHL via inhibiting both EGFR and CDK2 signaling pathways.

**Fig. 8.**
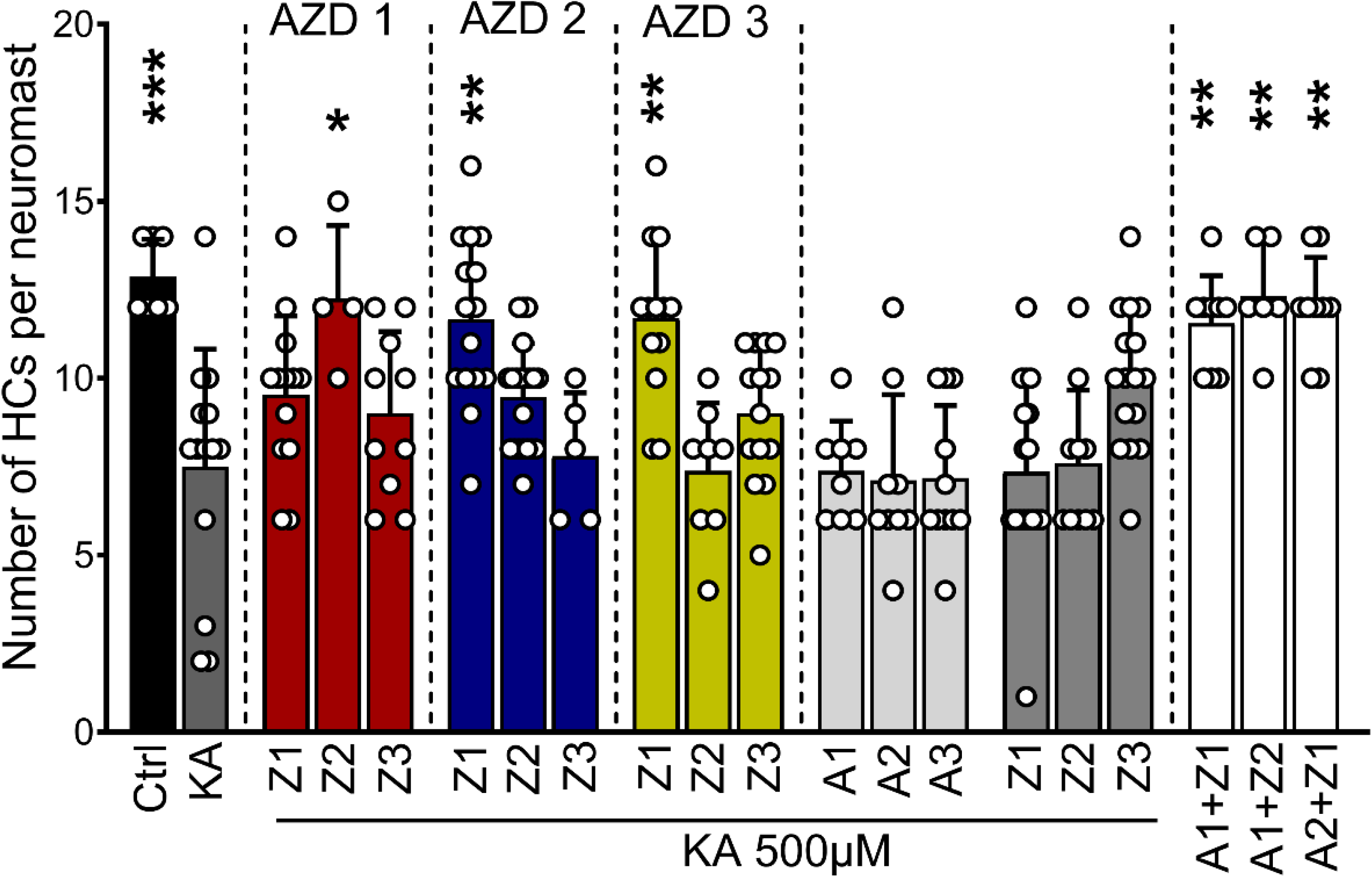
Synergistic otoprotection between zorifertinib and AZD5438 against excitotoxicity in zebrafish neuromasts. 5dpf zebrafish were pre incubated with KA (500µM) or regular fish water for 1 hour and then incubated for 2 hours with a combination of AZD5438 (CDK2 inhibitor) and zorifertinib (EGFR inhibitor) or with the individual compounds. Results are expressed as number of hair cells per neuromast ± SD. *P<0.05, **P<0.01 and ***P<0.001 versus KA alone (One-way ANOVA). A1: AZD5438 50nM, A2: AZD5438 1nM, A3: AZD5438 10pM, Z1: Zorifertinib 50nM, Z2: Zorifertinib 1nM, Z3: Zorifertinib 10pM.

## DISCUSSION

Noise-induced hearing loss (NIHL) is one of the most common causes of hearing impairment among military personnel, veterans and civilians (*1–4*). Despite the advances in the understanding of the underlying pathology associated with hearing loss, there are no drugs approved by the Food and Drug Administration (FDA) to prevent or treat NIHL. There remains an urgent unmet medical need for drugs to treat hearing loss (*60*). However, suitable high-throughput screening assays are currently unavailable for NIHL drug discovery. Animal model-based phenotypic screening for NIHL is relatively low-throughput, expensive and is often faced with difficulty in target deconvolution. To overcome these challenges, in this study, we utilized a novel strategy for the discovery of otoprotective compounds using cochlear transcriptomes associated with noise exposure and comparing those to a database of drug induced transcriptomic changes in cell lines. We identified 22 biological pathways and 64 drugs that have potential as otoprotectants to reverse the transcriptomic changes associated with noise exposure. We successfully validated in animal models EGFR inhibitors as promising otoprotective drugs for the treatment of NIHL.

### In silico screens for therapeutic drugs and biological pathways for hearing loss

Accelerating drug discovery for preventable conditions like NIHL is of paramount importance than ever before. In the past two decades, the utilization of transcriptional profiles and other omics data has seen a significant rise in the areas of pharmaceutical design, determination of drug mechanisms, and structure-activity relationship (SAR) evaluations (*61–63*). Different approaches are available for drug repurposing and one of the most commonly used strategies is transcriptomic signature comparison. Ever since the CMap (*30*) was first introduced, this method has been utilized in pharmacological research to establish connections between disease states and drugs (*64–66*). This method has also been employed by other scientists in their quest to discover potential new treatments for various types of cancer, along with rare conditions like Hirschsprung disorder (HD) and most recently for COVID-19 (*67–72*).

The overarching hypothesis of our study was that compounds able to revert the expression of genes associated with noise damage may be able to revert or prevent NIHL. In this study, we demonstrated that a data-driven analysis based on CMap represents a suitable approach for identifying new candidate drugs for NIHL. Our CMap analysis identified 22 biological pathways and 64 small molecule candidates including EGFR inhibitors as novel classes of drugs that may be developed further for the treatment of NIHL. EGFR signaling is novel in NIHL and additional downstream targets can provide additional NIHL drug candidates and biomarkers for NIHL pharmacokinetic analysis. Combinatory treatments of drugs targeting EGFR and CDK2 signaling pathways are more effective in otoprotection than individual drug treatment in the zebrafish model here, a result that corroborates with our recent studies using a combination of Braf and CDK2 inhibitors in preventing NIHL and cisplatin-indued hearing loss (CIHL) in mice (*22*).

### Repurposing FDA-approved drugs for preventing NIHL

Many benchmark candidates (sodium thiosulfate, ebselen, N-acetylcysteine, and D methionine originally chosen as antioxidants against neurodegeneration) have shown some promise in pre-clinical and clinical trials against NIHL (*73*); however, to date, no drugs have been approved by the FDA for clinical use to prevent any forms of NIHL. Repurposing FDA-approved drugs offers many advantages over developing new chemical entities (NCEs) (*22, 25, 27, 74, 75*). The first and most significant is that the safety and pharmaco-kinetics/dynamics (PK/PD) profiles of FDA-approved drugs are well defined in their respective dosing and formulation requirements in both pre-clinical and clinical studies. With such data, even drugs conventionally considered to have undesirable safety profiles may be repurposed at tolerated lower dosages for otoprotection. Secondly, it is a fast and cost-effective path to clinics. Based on FDA data since 2003, the number of approved repurposed drugs has surpassed that of NCEs (*76*). The average length of time for drug repurposing is 6-8 years (compared to 14 years for regular NCEs) and the average cost is ∼40-60% of the regular $2 billion price tag for a NCE (*77*). Repurposing drugs can also be further expedited for orphan diseases (e.g., pediatric cisplatin-induced hearing loss).

### EGFR inhibitors as candidate drugs for clinical trials on hearing loss

We selected EGFR inhibitors as our top otoprotectant candidates against NIHL from our noise transcriptomic LINCS analysis. In the enrichment analyses, afatinib was the top-ranking drug that was included in multiple pathways suggested to be involved in the pathogenesis of NIHL. Afatinib is an FDA-approved second-generation, irreversible, dual EGFR and Her2 inhibitor, with an IC_50_ of 0.5 and 14 nM, respectively, and with excellent kinase selectivity (*78*). It is FDA approved as the first-line treatment for patients with advanced/metastatic non-small cell lung carcinoma (NSCLC) carrying EGFR mutations (*79*). Additional generations of EGFR inhibitors have been recently characterized with even better specificity, potency, pharmacokinetics and pharmacodynamics (PK/PD) properties, bioavailability and blood brain barrier (BBB) permeability (**Fig. 1C**). We selected and tested the otoprotective effect of zorifertinib, a 4^th^ generation inhibitor that has excellent BBB permeability and bioavailability (*80, 81*). Zorifertinib is currently in clinical development for the treatment of NSCLC with CNS metastases (*82*). Both afatinib and zorifertinib plus two other EGFR inhibitors (osimertinib and dacomitinib) showed individually excellent protection, in multi-doses, against glutamate excitotoxicity that mimics noise damage in the zebrafish lateral line neuromast model (**Fig. 3**). Furthermore, we have shown that combinatorial treatment with zorifertinib and a specific, potent CDK2 inhibitor AZD5438, exhibits synergistic otoprotection in the zebrafish model (**Fig. 8**). We tested protection against NIHL using a permanent threshold shift noise exposure paradigm in a mouse model. Pretreatment with afatinib or zorifertinib provided 20-30 dB of otoprotection at all tested frequencies (**Fig. 4**). Exposure to octave band noise in mice reportedly produces maximum damage in cochlear regions that are one-half to two octaves higher than the exposure stimulus (*83*). In this study, significant otoprotection was observed even in the 64 kHz regions. To our knowledge, this is one of the first studies that has demonstrated otoprotection in the high-frequency cochlear region. Our mass spectrometry results provide direct evidence that zorifertinib crosses the blood-labyrinth barrier (BLB) (**Fig. 7**). Zorifertinib displayed excellent pharmacokinetic properties that are desirable for BBB/BLB penetrating drugs (*84*). Most importantly, our Pax2-Cre; EGFR conditional knockout (cKO) mouse (**Fig. 5**) and EGF knockdown zebrafish (**Fig. 3**) studies exhibited otoprotection against NIHL, similar to afatinib and zorifertinib; EGF knockdown plus afatinib in zebrafish did not show additional protection to that by either afatinib or EGF knockdown alone. Together these results confirmed that EGFR plays a key role in NIHL and mediates EGFR inhibitors’ otoprotection in vivo. Our results further demonstrate that EGFR inhibitors are an excellent class of drug candidates for NIHL and thus ready for advancing to clinical trials on hearing loss. Additional newer EGFR inhibitors with better specificity and PK/PD characters than afatinib and zorifertinib may therefore have potential as otoprotectants against NIHL and other forms of hearing loss.

### Mechanisms of action by afatinib and zorifertinib for otoprotection against NIHL

It has been well documented that afatinib and zorifertinib are potent EGFR inhibitors; however, it is unclear whether they act in similar mechanisms in cochleae against NIHL. To address this, we first confirmed that phosphorylation of AKT and ERK (two main downstream targets of EGFR) are indeed activated by noise exposure but mitigated by zorifertinib treatment. Importantly, we further performed an unbiased kinome analysis of mouse cochlear lysates using a novel kinase peptide array (*85, 86*). This experiment revealed multiple kinase-signaling pathways that are altered by noise exposure and ameliorated by zorifertinib in mouse models of NIHL (**Fig. 6E-I; Supplemental Figs. 34-14**), consistent with our Western results (**Fig. 6A-D**). Our Western and kinome results are also consistent with previous publications on noise-induced activation of several signaling pathways (AKT and ERK; (*56–59*)). Moreover, our kinome results are consistent with a recent study using proteomics under similar noise exposure in mice (*87*). Compared to previous studies, our kinome analysis uncovers a wider range of signaling pathways including more than 340 kinase targets/reporters, most of which are active in cochleae in response to noise exposure. It will be interesting to further examine lower doses of zorifertinib as well as various levels and durations of noise exposure to accurately profile the specific kinase pathways that are at works for NIHL.

Our results further support that inhibition of EGFR has effects at least similar to (if not more potent than) inhibition of individual downstream targets and that combinations of EGFR and additional inhibitors of downstream or synergistic pathways (i.e., AZD5438 for CDK2) are more effective than individual inhibitors in protecting against NIHL. Moreover, it remains unclear which cell types mediate EGFR effects during noise exposure. Given EGF ligands, EGFRs and downstream targets are expressed in both supporting cells, hair cells, and spiral ganglia (**Fig. 2**), it remains to be further studied how EGFR signaling mediates NIHL at cellular levels. It would be interesting to compare changes of kinome signaling pathways under other ototoxic insults (cisplatin, antibiotics and aging) so that both common and specific cochlear mechanisms of action can be validated, and corresponding therapeutic strategies can be developed.

An increasing number of recent studies have surprisingly revealed that signaling pathways normally controlling cell proliferation in cancers are also involved in stress-induced cell death in post-mitotic, wild-type cells (*22, 25, 27, 74, 75*). For example, CDK2 and Braf inhibition protected against ototoxic insults in postmitotic cochlear cells as we have demonstrated here on EGFR inhibition (*22*). These studies together suggest that pharmaceutical inhibition of cell proliferation pathways (i.e., EGFR, CDK, and Braf) may have similar protective effects against stresses in nervous and other post-mitotic systems.

### Limitations

Our in silico screens are based on transcriptomic changes of mostly drug-treated cancer cell lines in CMap that heavily reflect drug effects on cell proliferation while additional pathways are likely involved in stress responses that remain to be further identified. Additionally, our screens assumed that pathway and drug hits are directly involved in reversing the damage caused by noise rather than reversing possible protective pathways. From the list of 22 biological pathways and 64 drug candidates in our in-silico screens, many have been tested as otoprotective in various in vitro or in vivo models; however, we have only tested one top pathway (EGFR) and its inhibitors. It remains to be further tested if other pathways and drugs work under similar NIHL conditions. Moreover, the doses at which zorifertinib and afatinib were tested could induce multiple non specific pathways as evidenced in our kinome panel results; it is desirable to test lower doses of drugs for specific pathways involved. Finally, we only tested one noise exposure condition and one drug regimen in mice; it is important to test additional levels, durations of noise and drug regimens to further elucidate the full potential of EGFR inhibitors in protecting against various forms of NIHL.

## MATERIALS AND METHODS

### Ethics Statement

Care and use of animals followed the guidelines in the NIH Guide for the Care and Use of Laboratory Animals. All animal procedures were approved by the Institutional Animal Care and Use Committee of Creighton University. All efforts were made to minimize pain.

### Materials

Afatinib dimaleate was purchased from Cayman Chemical, USA. Zorifertinib (AZD3759) and AZD5438 were purchased from MedChemExpress, USA. Antibodies used included: C terminal binding protein-2 (mouse anti-CtBP2; BD Transduction Labs, used at 1:200), myosin-VIIA (rabbit anti-myosin-VIIA; Proteus Biosciences, used at 1:250), anti-otoferlin (HCS-1, DSHB 1:500), anti-GFP (NB100-1614, Novus Biologicals 1:500), total AKT and ERK (Cell Signaling 9272S and 4695S, respectively), phospho forms (AKT-S473 and ERK1/2-T202/Y204, Cell Signaling) and β-actin (Sigma A3854).

### Drug identification using LINCS Query

Microarray and RNA-seq transcriptomes from cochlea following noise exposure that are available in the public Gene Expression Omnibus (GEO) database were analyzed using the National Center for Biotechnology Information (NCBI)’s GEO2R tool (https://www.ncbi.nlm.nih.gov/geo/geo2r/) to identify differentially expressed genes between the two groups. Genes with an absolute log-fold change greater than 1 were downloaded from each study and analyzed with LINCS databases to identify compounds inducing similar gene expression profiles in various cell lines. The LINCS analysis relies on a subset of the 1,319,138 genetic profiles originally compiled in the L1000 compendium. For each profile, an overlap score between 0-1 was given, indicating the fraction of genes that either mimic or reverse the gene set input. With over 100 identified compounds of interest, we further narrowed down the results of our screen by selecting those compounds with an overlap score >0.1, indicating at least a 10% overlap between the small molecule perturbation from the databases and our gene expression profile.

Three comparison groups were created from the Gratton, et al. (*40*) DNA microarray data set. The first experimental group compared transcriptomes of the 129X1/SvJ mouse without noise exposure to the B6.CAST mouse without noise exposure, This group served as our control group (N-/-) and identified DEGs that may have conferred resistance to NIHL in the 129X1/SvJ mouse prior to noise exposure. The second experimental group compares the transcriptomes of the 129X1/SvJ noise treated mouse to the 129X1/SvJ control mouse and is referred to as (129 N+/-). The purpose of this group is to determine which genes may be involved in hearing protection for the 129X1/SvJ following noise exposure. The third experimental group compares the transcriptomes of the B6.CAST noise-treated mouse to B6.CAST control mouse and is referred to as (B6 N+/-). The purpose of this group is to determine which genes may be involved in noise trauma ototoxicity after noise exposure.

12,488 differentially expressed genes (DEGs) were identified for each of the three experiment groups using GEO2R and were ranked based on p-value and log-fold change. DEGs with a log fold change greater than 2 and p-value less than 0.05 were considered differentially expressed genes of interest. Using this threshold, 92 upregulated genes and 146 downregulated genes were found for the N-/- group, 138 upregulated genes and 24 downregulated genes were found for the 129 N+/- group, and 109 upregulated genes and 41 downregulated genes were found for the B6 N+/- group. DEGs of interest were used for Gene Ontology pathway analysis using the ShinyGO enrichment analysis program (http://bioinformatics.sdstate.edu/go76/). This program identified 30 biological pathways for each experimental group. Pathways were ranked based on enrichment FDR value and the program identified which of the input genes were significant for each biological pathway.

L1000CDS^2^ analyses were performed using the DEGs from each biological pathway with at least 3 upregulated and 3 downregulated DEGs of interest. Each L1000CDS^2^ analysis reveals 50 drug perturbations that mimic or reverse the input transcriptome and are ranked based on overlap score. In total, 65 L1000CDS^2^ analyses were performed; 27 analyses from the N-/- Mimic group, 13 analyses from the 129 N+/- Mimic group, and 25 from the B6 N+/- Reverse group. Therefore, 1,350 drug perturbations were found that mimic the N-/- DEGs, 650 drug perturbations that mimic the 129 N+/- DEGs, and 1,250 drug perturbations that reverse the B6 N+/- DEGs. Drug perturbations with the highest overlap scores were considered significant and filtered the list of significant drug perturbations down to 189 significant drug perturbations from the N-/- Mimic group, 53 significant drug perturbations from 129 N+/- Mimic group, and 173 significant drug perturbations from the B6 N+/- Reverse group. Drug perturbations were further filtered by targeting which drugs target multiple pathways. In total, 83 drug perturbations were found to target multiple pathways between the three experimental groups.

Two experimental groups were created from the Maeda et al. (*41*) bulk RNA-seq data set. The transcriptome of the C57B6 mouse without noise exposure was compared to the C57B6 mouse after noise exposure and identified 939 DEGs. DEGs with a log fold change greater than 2 were considered significant. Of the 939 DEGs, 51 significant upregulated genes and 222 significant downregulated genes were identified. In addition, the Maeda group examined significant DEGs that encode for transcription factors and identified 9 significant upregulated genes and 16 significant downregulated genes. The first experimental group used all 51 upregulated genes and 222 downregulated genes as input to a L1000CDS^2^ analysis. The second experimental group used the 9 upregulated genes and 16 downregulated genes that encode for transcription factors as input to a L1000CDS^2^ analysis. The purpose of these two experiments was to identify drug perturbations that would reverse the transcriptome of the noise-exposed C57B6 mouse. Each of these experiments identified 50 drug perturbations that were ranked based on overlap score for a total of 100 drug perturbations. Of these 100 drug perturbations identified, 43 drug perturbations of these were also identified from the Gratton et al. data set.

For the third dataset from Milon et al.(*42*), DEGs for outer hair cells and supporting cells (6 and 24 hrs), marginal, intermediate, basal cells, fibrocytes from the cochlear lateral wall, and the spiral ganglion neurons were used as input for L1000CDS^2^ queries. DEGs with a log fold change greater than 1.2 were considered to be significant. For each cell type, the top 50 drugs ranked according to the cosine distance score were compiled. From the list of 500 drug perturbations, drugs, and mechanism of action classes were compared, and a consensus list was prepared for the three datasets.

### Molecular docking

Structure modeling, docking and analysis were done using the YASARA package (*88*). For docking dacomitinib and zorifertinib to the EGFR, the crystal structure of EGFR kinase – afatinib complex was used (PDB id.4G5J). The missing residues 747-756 (LREATSPKAN) from the X-ray structure were inserted using the loop modeling option of YASARA. The completed structure was solvated with water molecules in a rectangular box so that the distance between protein and the box edge was 10 Å. The structure of the solvated system was energy-minimized using the AMBER ff14SB force field (*89*) and then it was subjected to 1 ns molecular dynamics (MD) simulation at 300 K temperature and 1 atm pressure. Using the last frame of the MD trajectory, afatinib was removed from the complex and the structures of dacomitinib and zorifertinib, obtained from https://pubchem.ncbi.nlm.nih.gov, were docked to the receptor using AutoDock VINA (*90*) in YASARA. Molecular contact surface area and contact area color determined by the ESPPBS method, which includes the implicit water effects.

### Animals and drug administration

#### Zebrafish

Danio rerio experimental larvae were obtained by pair mating of adult fish maintained at Creighton University by standard methods approved by the Institutional Animal Care and Use Committee. We used Tg(brn3c:mGFP) expressing a membrane-bound GFP in HCs. Experimental fish were maintained at 28.5°C in E3 water (5 mM NaCl, 0.17 mM KCl, 0.33 mM CaCl_2_, and 0.33 nM MgSO_4_, pH 7.2). Animals were cryoanaesthetized after drug treatment and prior to fixation. The neuromasts inspected, SO3 and O1-2, were part of the cranial system.

#### Mice

This study used 7 to 10-week-old FVB/NJ mice obtained from The Jackson Laboratory (Bar Harbor, ME, USA), with an equal number of males and females across the experiments. Conditional EGFR knockout mice were generated by crossing floxed EGFR mouse strain – (Egfr^tm1Dwt^ (*91*), Stock# 031765-UNC, Mutant Mouse Resource and Research Center) with Pax2 Cre strain (*92*). The mice were on mixed background (CD1, CBA/CaJ, C57BL/6).

### Zebrafish drug studies

The lateral-line neuromasts of zebrafish are a valuable system for testing protectivity of compounds against cisplatin toxicity in vivo, as their HCs are considered homologous to those in the mammalian inner ear and are readily accessible to drugs.

To test whether the drugs protect HCs from excitotoxic trauma, we employed a zebrafish model that mimics noise damage by exposing fish to the ionotropic glutamate receptor agonist, N-methyl-D-aspartate (NMDA) previously shown to cause progressive HC loss in zebrafish lateral-line organs. Briefly, 5-dpf larvae were preincubated with 300 µM NMDA for 50 minutes followed by 2 hrs of incubation with the drugs at 0.002 µM and 0.0183 µM.

Subsequently, animals were transferred to E3 water for 1 hour, fixed in 4% paraformaldehyde (PFA) overnight and immunostained for otoferlin and GFP. HCs were manually counted using a Zeiss AxioSkop 2 fluorescence microscope with a 40x oil objective. Compounds were then evaluated on efficiency and potency, with the top-rated compounds showing high protection at lower concentrations. For neuromast imaging, samples were analyzed under a Zeiss LSM 700 confocal microscope with an oil immersion objective of 63X (numerical aperture 1.4) and 1.3x digital zoom.

### Noise exposure

FVB/NJ mice (6 to 7-weeks old) were exposed to noise inside a double-walled soundproof booth (IAC Acoustics). Awake mice held individually in small open-walled cylindrical containers inside a reverberant chamber were exposed to octave band noise (8-16 kHz at 100 dB SPL) for 2 hours delivered via an exponential horn fitted on to a titanium horn driver (JBL 2426H) and driven by a power amplifier (Crown CDi1000) using RPvdsEx circuit and a RZ6 Multi-I/O processor (Tucker Davis Technologies, Alachua, FL). Before each session, the overall noise level was measured and calibrated at the center and four quadrants of the reverberant chamber using a calibrated ¼” microphone (PCB 377C10). All noise exposures were from noon to 2PM to minimize differences in circadian variation in sensitivity to noise. Noise exposure level was based on our previously published work (*27*) showing 8-16 kHz octave band noise at 100 dB for 2 hours causes 20 dB ABR threshold shift and IHC synaptic ribbon loss.

### Auditory brainstem response (ABR)

Mice were anesthetized with avertin (5 mg/10 g body weight, IP) and placed on a homeothermic heating pad in conjunction with a rectal temperature probe. Subcutaneous electrodes were inserted at the vertex (reference), posterior to the pinna (active), and over the hip (ground). The acoustical stimulus generation, ABR wave acquisition, equipment control, and data management were performed using National Instruments PXI system with 6221 and 4461 modules and the EPL Cochlear Function Test Suite (CFTS). Stimuli were presented by a TDT MF1 driver in an open field configuration placed 10 cm from the left ear of each animal. Tone-pip stimuli of 5 ms duration at half-octave frequency intervals from 4.0-64 kHz were presented at 21/s. At each frequency, an intensity series was presented from 10-100 dB SPL in 5 dB incremental steps. 256 responses of alternating stimulus polarity were collected and averaged for each stimulus level. Evoked-response signal was amplified 10,000x (Grass P5-11 bio amplifier) and band pass filtered (0.3-3 kHz) before digitization. ABR threshold was determined as the lowest stimulus level that produced a detectable ABR waveform (wave-1 to 5) that could be visualized. ABR threshold shifts were calculated by subtraction of the pre-exposure thresholds from the post exposure thresholds.

### Distortion product otoacoustic emissions (DPOAEs)

The primary tones f1 and f2 were generated and shaped using EPL CFTS software and NI PXI system. The two primary tones were presented using two TDT MF1 speakers in closed field configurations. The primary tones were delivered using a custom probe insert attached a low-noise ER10B+ microphone (Etymotic Research, USA). DPOAEs were recorded in the form of level/frequency functions; f2/f1 was fixed at 1.2, with the level of the f2 (L2)10 dB less than the f1 level (L1). The f2 stimuli were presented at 5.6 – 32 kHz at half-octave intervals. At each f2 frequency, L2 was varied between 65 and 5 dB SPL at 10 dB steps. The 2f1-f2 DPOAE amplitude and surrounding noise floor were extracted by offline analysis. DPOAE threshold was defined as the L1 level that produced emission at 2f1-f2 with emission amplitude of 0 dB SPL. The average noise-floor was -25 dB SPL across frequencies.

### Sample preparation and immunofluorescence labeling

After the final ABR and DPOAE measurements, the mice were transcardially perfused with 4% paraformaldehyde in 0.1 M phosphate buffer. After perfusion, cochleae were isolated and post-fixed in 4% PFA in 0.1 M phosphate buffer for 2 hrs at room temperature. After fixation, the cochleae were decalcified in 120 mM EDTA for 2-3 days. Each cochlea was microdissected into three pieces and a cochlear frequency map was computed based on 3D reconstruction of the sensory epithelium for outer hair cells (OHCs) and presynaptic ribbon counts. Dissected pieces were permeabilized using 0.02 % Triton X-100 for 15 min, washed three times in PBS and preincubated for 1 hr in blocking buffer (10% normal goat serum) at room temperature. Cochlear pieces were incubated with CtBP2 or myosin-VIIa, with matching secondary antibodies (Alexa Fluor 488, 546; Life Technologies, USA). DAPI was used for nuclear staining (DAPI, Thermo Fisher, 1:1000). Stained cochlear pieces were mounted on slides with Fluoromount-G medium (SouthernBiotech, USA) and cover-slipped.

### Quantification of synaptic ribbons

Following the frequency map computation, cochlear structures were located to relevant frequency regions. Using a confocal microscope (Zeiss LSM 700), OHC and IHC zones were both imaged with a 63x, numerical aperture 1.4 with 1.0x digital zoom. For IHC ribbon synapse quantification, 3D (x-y-z-axis) images were scanned with the 1.3x digital zoom at 63x. Each immunostained presynaptic CtBP2 puncta was counted as a ribbon synapse. Synaptic ribbons of ten consecutive IHCs distributed within the 16-22 kHz frequency region were counted. The CtBP2 (presynaptic) puncta were counted using Imaris 9.5 (Oxford Instruments, Abingdon, UK) using the approach described by Fogarty et al. (*93*). The average spot diameter was set to 0.45 μm, and only CtBP2 puncta found within the surface of the IHC were included. The synaptic ribbons in the normal cochleae were also calculated using the same method to serve as control comparison samples.

### Western blotting

For the immunoblot studies, animals were administered afatinib, zorifertinib or the vehicle 1 day before and immediately after the noise-exposure. Animals were sacrificed either 30 minutes after the noise-exposure and the cochleae isolated and processed in lysis buffer (RIPA, ThermoFisher, 89901) containing protease inhibitors. Fifteen to thirty micrograms of protein were used for the immunoblot experiments. Membranes were blocked with 3% of milk blocking solution and incubated with the primary antibodies overnight at 4°C. After several washes and the incubation with the secondary antibody, membranes were developed using a ChemiBlot system (Bio-Rad). Membranes were stripped and reprobed for the phosphorylated forms. β-actin was used as the loading control.

### Kinome analyses

#### Identification of significant differential AKT and ERK family kinase activity

24 AKT and 25 ERK kinase family putative target peptides were identified by the KRSA package (*94*). Log2-fold changes in peptide activity were calculated by comparing Noise, Zorifertinib, or Noise + Zorifertinib to control. For each peptide, an average of triplicates was used per condition. Two-way ANOVA was used to identify significant differences between groups (****: P<0.0001, ***: P<0.001), with results presented as mean ± standard deviation.

### Methods Omnibus for the PamGene Kinome Array

#### Overview

The PamGene platform is a well-established, highly cited, microarray technology for multiplex kinase activity profiling (*85, 86, 95, 96*).

#### Hardware

Pamstation12 and PamChip4. The PamGene12 Kinome Array is a peptide array based platform that facilitates the unbiased detection of kinase activity by serine/threonine (STK) or tyrosine (PTK) kinases (*85, 86, 95-98*). The STK and PTK PamGene12 chips have 144 and 196 reporter peptides, respectively. Each spot has approximately 300,000 copies of the same peptide printed on it. Phosphorylation is detected in real time. Fluorescent antibodies are applied against phosphorylated residues; fluorescent intensity is a proxy for the extent of reporter peptide phosphorylation (**Supplemental Fig. 4**). Altered kinase activity can be directly measured. For example, phosphorylation by PKA on the STK chip is concordant with its activity in solution (*95*). Each peptide chip has 4-wells, and three chips can be run at the same time. Thus, there are up to 12 samples for each “run” on the array.

#### Chip Coverage

Of the about 500 kinases in the human genome (*99, 100*), 245/376 (65%) Ser/Thr and 89/93 (96%) Tyr kinases can be mapped to the STK and PTK chips, respectively. The chips also map about 18/21 (86%) dual specificity kinases, covering about 72% of the entire kinome. The STK chip covers similar amounts of low (52%), medium (65%), and high (65%) abundance protein kinases in neurons (based on Brainseq neuron database (https://www.brainrnaseq.org/)) and has sensitivity for detection into the picogram range (unpublished data from Pamgene) for many kinases.

### Kinome Array Protocols

#### Data Generation

Samples are prepared according to the protocols provided by PamGene Corp (https://pamgene.com/ps12/). The catalytic activity and stability of kinases are controlled by the addition of protease and phosphatase inhibitors. Peptide phosphorylation is monitored during the incubation with assay mixture, by taking images every 5 min for 60 min at exposure lengths of 5 msec, 25 msec and 100 msec, allowing real time recording of the reaction kinetics. Various internal control tests have been performed by PamGene International to ensure the sensitivity of the assay. Chip-to-chip and run-to-run technical variation (coefficient of variability (CV)) is <9% and <15%, respectfully. To account for technical variation between runs, an internal control sample may be added to account for between run variability.

#### Preliminary Data Processing

The primary output from PamStation12 is images from the Evolve kinetic image capture software. These images are then pre-processed to quantify the activity at each peptide level using the PamGene’s BioNavigator software (https://pamgene.com/wp-content/uploads/2020/09/BioNavigator-User-Manual-vs2.3-2020.pdf). Before proceeding to activity analysis, all peptides that appear as inactive (Raw Signal <= 5) are removed from the analysis. The dynamic range of the raw signal intensities is typically 0 - 3,000. Linear regression slope of the signal intensity as function of exposure time is used to represent the peptide phosphorylation intensity for downstream comparative analyses, averaged across the biological replicates. This is done to increase the dynamic range of the measurements. The signal ratio between case and control samples is used to calculate fold change (FC) values. Peptides with a fold change of at least 15% (ie FC > 1.15 or FC < 0.85) are considered differentially phosphorylated for the purposes of using KRSA. This threshold was chosen based on previous reports that suggest small changes in kinase activity are sufficient to trigger biologically relevant changes (*97, 98, 101*). Peptides that had very low signal or an R^2^ of less than 0.90 during the corresponding linear regression are considered undetectable or non-linear in the post-wash phase and were excluded from subsequent analyses.

#### Assessment of Upstream Kinases

Peptides spotted on the array (and in general) may be phosphorylated by more than one kinase, and in many cases several different kinases. The use of two different types of chips, one for Ser/Thr kinases (STK) and one for Tyr kinases (STK) provides a starting point for assignment of kinases. There are 4 different software packages that may be deployed for assignment of upstream kinases. All of them rely, to varying extents, on publicly available mapping databases. Each has strengths and weaknesses, some of which are discussed below.

#### Upstream Kinase Analysis (UKA)

This package was developed by the Pamgene Corp (s’ Hertengobosch, Netherlands). UKA is integrated into the manufacturer’s BioNavigator software and their recommended method. This method relies on a curated database of kinase substrate interactions created by the PamGene Corp. It takes the raw output from the PamStation as input. It then filters low intensity peptides and scales the entire dataset to the range of 0-100. It then calculates a “kinase Score” for each kinase and reports the ones with the highest score. Advantages include 1) providing results for specific kinases (as opposed to families) and 2) a low false positive rate compared to other packages. One putative weakness is that it may be too stringent for discovery-based experiments.

#### Kinome Random Sampling Analyzer (KRSA)

The package was developed by the Cognitive Disorders Research Laboratory (CDRL) at the College of Medicine and Life Sciences (COMLS) University of Toledo, led by Dr. Robert McCullumsmith (*94*). The data generated from the kinome array experiment and the mapping of the PamChip file are used as input to the algorithm. Once selected, peptides are filtered out using advancement criteria, including the signal intensity at maximum exposure time and the R2 values of the linear regression of signal intensity as a function of exposure time. At the end of this step, a list of filtered peptides moves forward to the next step of the analysis.

#### Curation of the database of upstream kinases

KRSA relies on a curated database of upstream kinases for the peptides present on the array. Protein kinases predicted to act on phosphorylation sites within the array peptide sequences were identified using GPS 3.0 and Kinexus Phosphonet (Kinexus Bioinformatics) (*102–104*). These programs provide predictions for serine-threonine kinases targeting peptide sequences ordered by likelihood of binding. The union of the highest ranked 5 kinases in Kinexus and kinases with scores more than two times the prediction threshold in GPS 3.0 were considered predicted kinases for each peptide and used in KRSA analysis (*95*). This list was combined with kinases shown in the literature to act on the phosphorylation sites of the peptides via PhosphoELM (http://phospho.elm.eu.org) and PhosphoSite Plus (https://www.phosphosite.org).

### Presentation of the data in KRSA

#### Heatmaps

Heatmaps are generated from the signal intensity data. The selection of peptides is based on the quality control criteria explained above. The values on the heatmap are the slopes of the linear models of signal intensity as a function of the exposure time.

#### Violin Plots

Violin plots showcase the distribution of the signal intensity of significant peptides on a per-group basis.

#### Waterfall Plots

Waterfall plots are generated from the Z score values for each kinase. These values are generated on a chip-by-chip basis and then averaged across the three. The plot shows the distribution of these three points and a red dot showing the mean z score value.

#### Kinase Enrichment Analysis 3 (KEA3)

KEA3 is an upstream kinase assignment method developed by the Maayan laboratory (https://maayanlab.cloud/kea3/) that relies on the known kinase protein interactions and kinase substrate interaction data and associated co-expression and co-occurrence data (*105*). The KEA3 web server takes a set of phosphorylated proteins and their fold-change value as input and returns the putative upstream kinases using the database and looking for statistically significant over representation of kinases.

#### PTM Signature Enrichment Analysis (PTM-SEA)

PTM-SEA (*106, 107*) is an application developed by the Broad Institute to identify putative upstream kinases. PTM-SEA is a modified form of the single sample Gene Set Enrichment Analysis (ssGSEA) with the underlying database built on top of PTMSigDB (*106*). The software runs on the R Programming language. It takes in files in the Gene Cluster Text format which has the peptides and log fold change values in a specified format. The output from PTM-SEA is a list of putative upstream kinases that can phosphorylate each site.

#### Integration of Upstream Kinase Assignments Across Packages

All the tools above utilize different methods to assign upstream kinases. This necessitates the use of an integration system to identify consensus upstream kinases across datasets. For this purpose, we utilize the software Creedenzymatic (*108*). The Creedenzymatic analysis takes in the results from at least two of the 4 analyses mentioned above and then generates a consensus figure with kinases deconvolved and ranked based on their presence in the results.

### Perilymph sampling and quantitative mass spectrometry

FVB mice (6-8 weeks old) were administered zorifertinib (15 mg/kg) via oral gavage. Inner ear perilymph fluid was collected before and after (30 min, 1 hr, 2 hr, 4 hrs, and 6hrs) the drug treatment. 1 μl samples were collected from the posterior-most, extracranial portion of the posterior semi-circular canal (*109*) and diluted 50-fold with 0.1% formic acid in water. Samples were frozen for later analysis and quantified via our LC-MS/MS in the Mass Spectrometry Core. Mice with vehicle treatment only were used as negative controls. Samples were spiked with a standard amount of 1 μg/ml IS (crizotinib). 20 µl of the diluted sample were injected in a Vanquish UPLC through a Waters Acquity BEH 18C 1.7 µm x 2.1 µm x 50 mm column coupled to a Q exactive quadrupole mass spectrometer with electrospray ionization (ESI) interface. The column was maintained at 40°C. The mobile phase consisted of eluent A (0.1% formic acid in water) and eluent B (acetonitrile) at a flow rate of 0.2 ml/min with the following gradient: 0 to 1 mins – 5%B; 1 to 4 mins – 40%B; 4 to 6 mins – 50%B;.6 to 8 mins – 65%B, 8 to 10 mins 100%B; 10 to 22 mins – 5%B. ESI conditions were: spray voltage – 3.9 kV; Capillary temp. 320°C. The PRM inclusion list was 460.153 (zorifertinib) and 450.13 (IS), and detection was run in positive po. All acquisition and analysis of data was done using Xcalibur software (Thermo Fisher Scientific, MA, USA).

### Statistical analysis

All statistical analyses and graphical visualization were performed in GraphPad Prism v9.x (GraphPad, MA, USA). Comparisons between the treatment groups for ABR and DPOAEs were performed using two-way ANOVA followed by Holm-Šidák post hoc test. A paired student’s t-test was used for comparison of the CtBP2 puncta between experimental groups. ABR/DPOAE thresholds, and CtBP2 puncta counts were determined by an independent observer who was blinded to the treatment condition. For zebrafish quantification studies one-way ANOVA was performed followed by Dunnet post-hoc test. Statistical significance was set at p-value ≤0.05. Unless stated otherwise, the results are expressed as mean ± SEM.

## Acknowledgement

This work was supported in part by NIHR01DC015010, NIHR01DC015444, USAMRMCRH170030/200079, ONR-N00014-18-1-2507, State of Nebraska LB692 (289325) and Stem Cell Subaward (3782) to JZ, USAMRMCRH190050 (to MZ) and the Bellucci Family Foundation Awards (to MZ and SV), 1P20GM139762 to Dr. Richard J. Bellucci Translational Hearing Center at Creighton University, MH107487, MH121102, and AG057598 to REM. The Pax2-Cre strain was gifted by Sonia Rocha-Sanchez, Creighton University.

---

**Supplemental Fig. 1.**
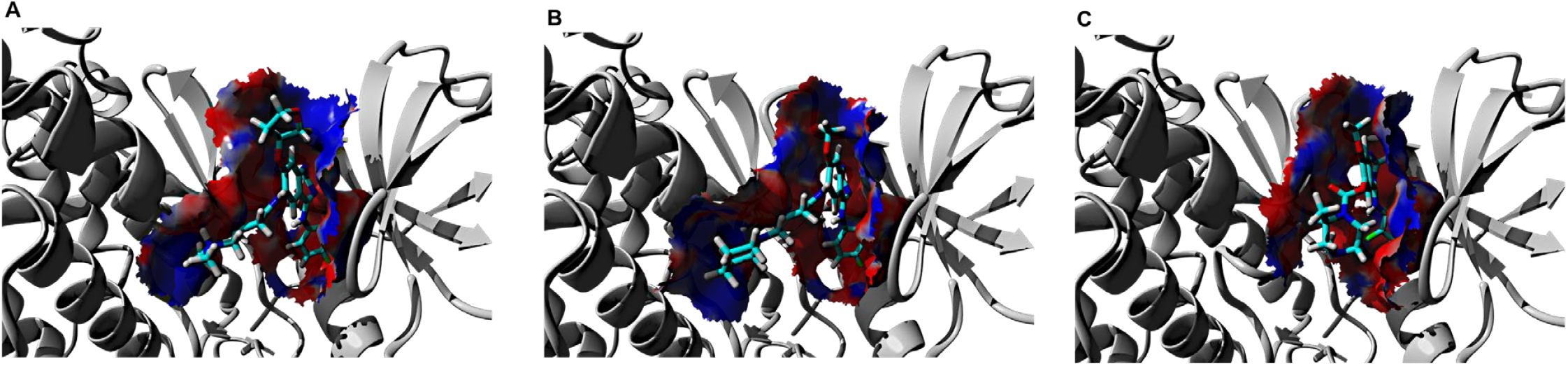
EGFR kinase inhibitors docked to the receptor binding site. **A**, Afatinib; **B,** Dacomitinib; **C** Zorifertinib. Cartoon representation of EGFR is in grey color, inhibitor molecules are in stick representation. Interacting molecular surface areas between the EGFR and inhibitors are colored by the electrostatic potential between the protein and ligand; red and blue colors represent the negative and positive potential, respectively. The main structural differences among these inhibitors appear at the crotonamide side chain which can be correlated to EGFR signaling differences.

**Supplemental Fig. 2.**
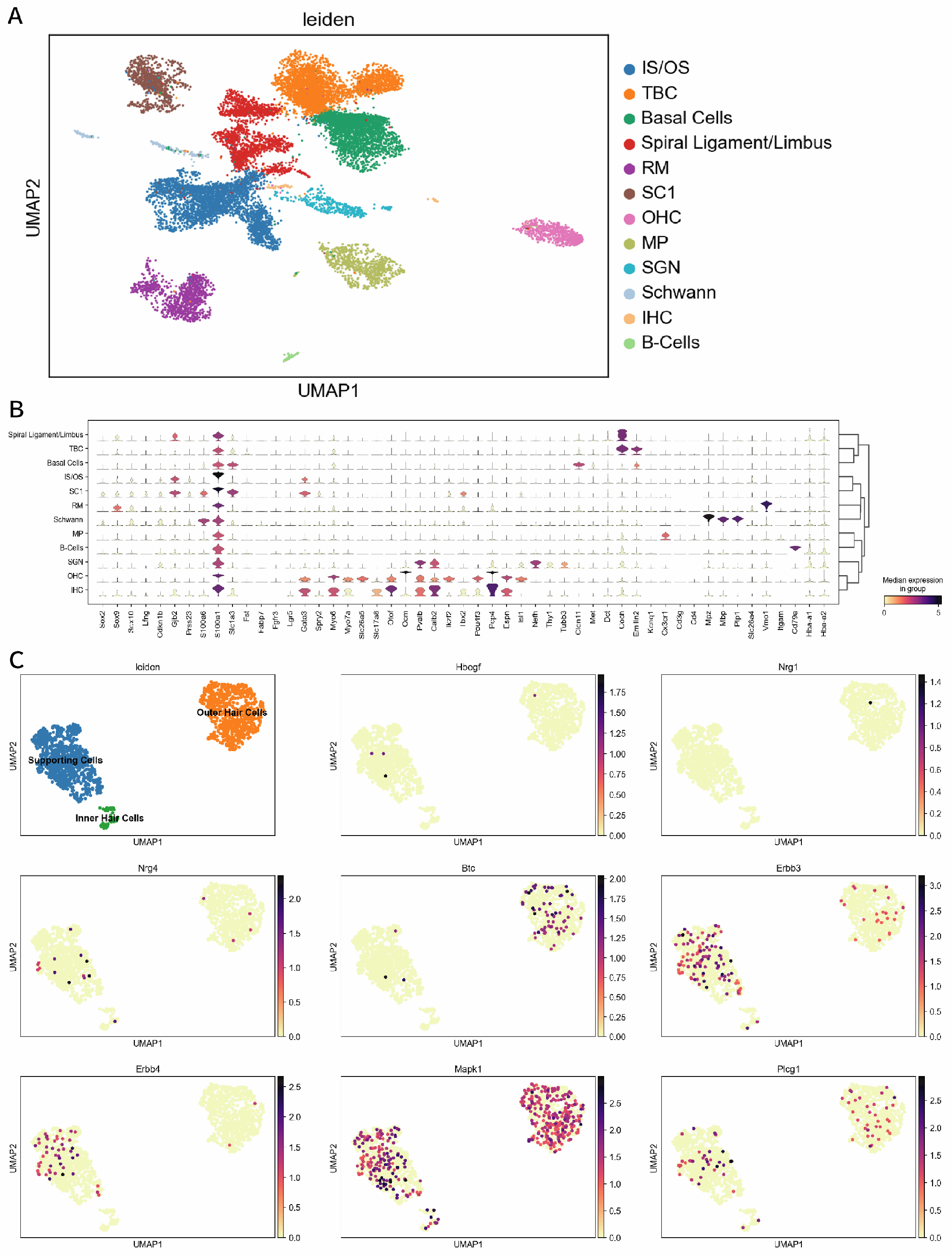
scRNA-seq analysis of EGFR signaling in adult mouse cochleae. (**A**) UMAP plot of the whole P28 C57/Bl6 cochlea dataset. (**B**) Stacked violin plot showing expression levels of marker genes ion all clusters. (**C**) UMAP plots showing Leiden clustering (for reference), and distribution and expression of ligands (Hbegf, Nrg1, Nrg4, Btc), receptors (Erbb3, Erbb4), and downstream targets (ERK/Mapk1, Plcg1) in supporting cell, outer hair cell, and inner hair cell clusters. IS/OS = Inner/Outer Sulcus, TBC = Tympanic Border Cells, RM = Reissner’s Membrane, SC1 = Supporting Cells, OHC = Outer Hair Cells, MP = Macrophages, SGN = Spiral Ganglion Neurons, IHC = Inner Hair Cells.

**Supplemental Fig. 3.**
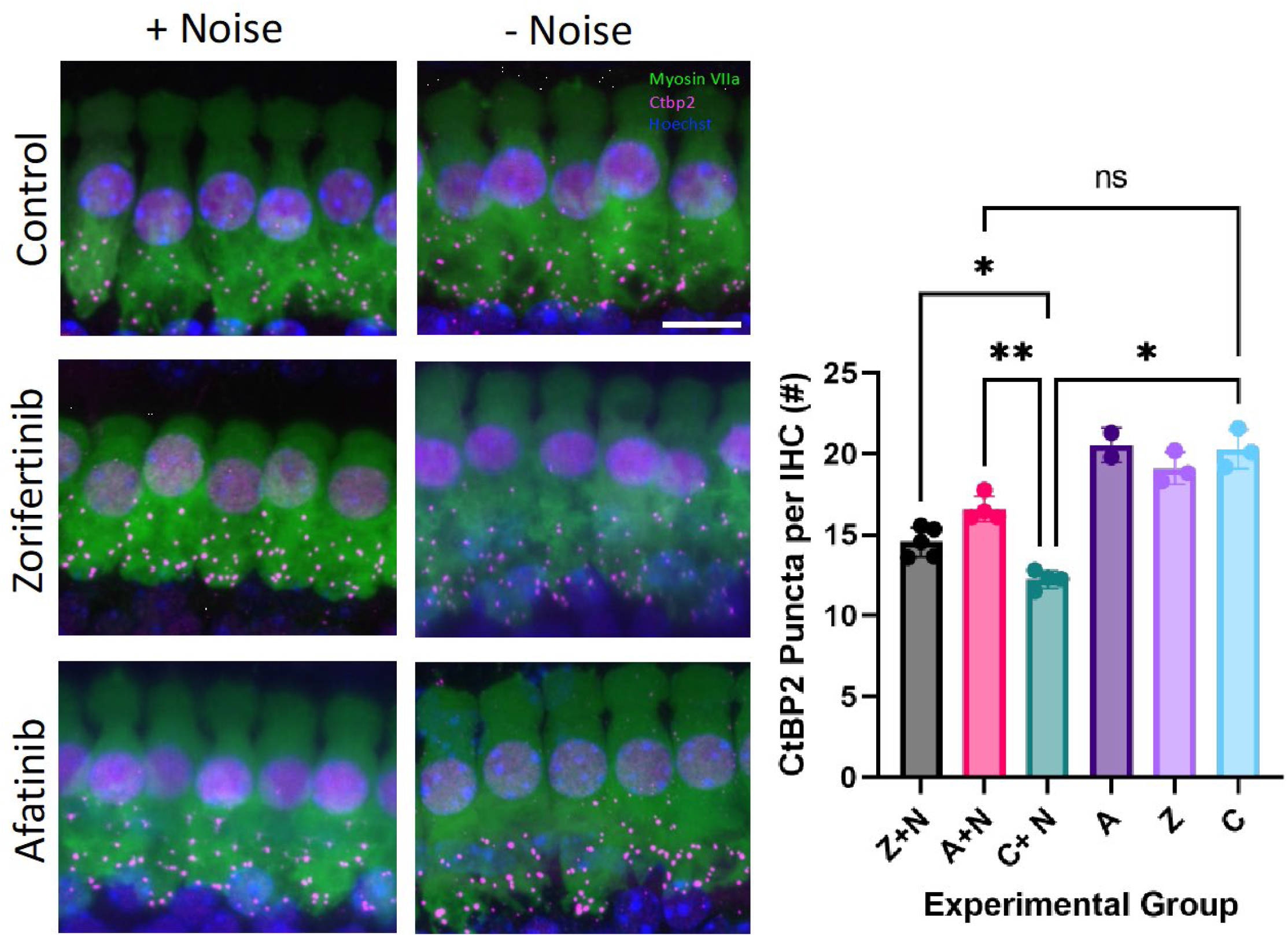
Afatinib and zorifertinib protect against noise-induced cochlear synaptopathy in mice. (**A**) Representative maximum intensity projections of inner hair cells (IHCs) in the 16-22 kHz region of the cochlea following drug treatment and with (left) or without noise (right). Hair cells were labeled using myosin-VIIa, presynaptic puncta were labeled using CtBP2, and nuclei were counterstained with Hoechst. (**B**) Zorifertinib and Afatinib protect against noise induced cochlear synaptopathy, resulting in less CtBP2 puncta loss with drug + noise than control + noise. The number of CtBP2 puncta per inner hair cell (IHC) is expressed as mean +/- SD; n=2- 5 animals per group. Each dot (n) represents one animal and the average CtBP2 puncta across ten IHCs from two cochleae. *P<0.05. **P<0.01, ***P<0.001; Welch’s ANOVA. Z: Zorifertinib; A: Afatinib; C: Control; N: Noise.

**Supplemental Fig. 4.**
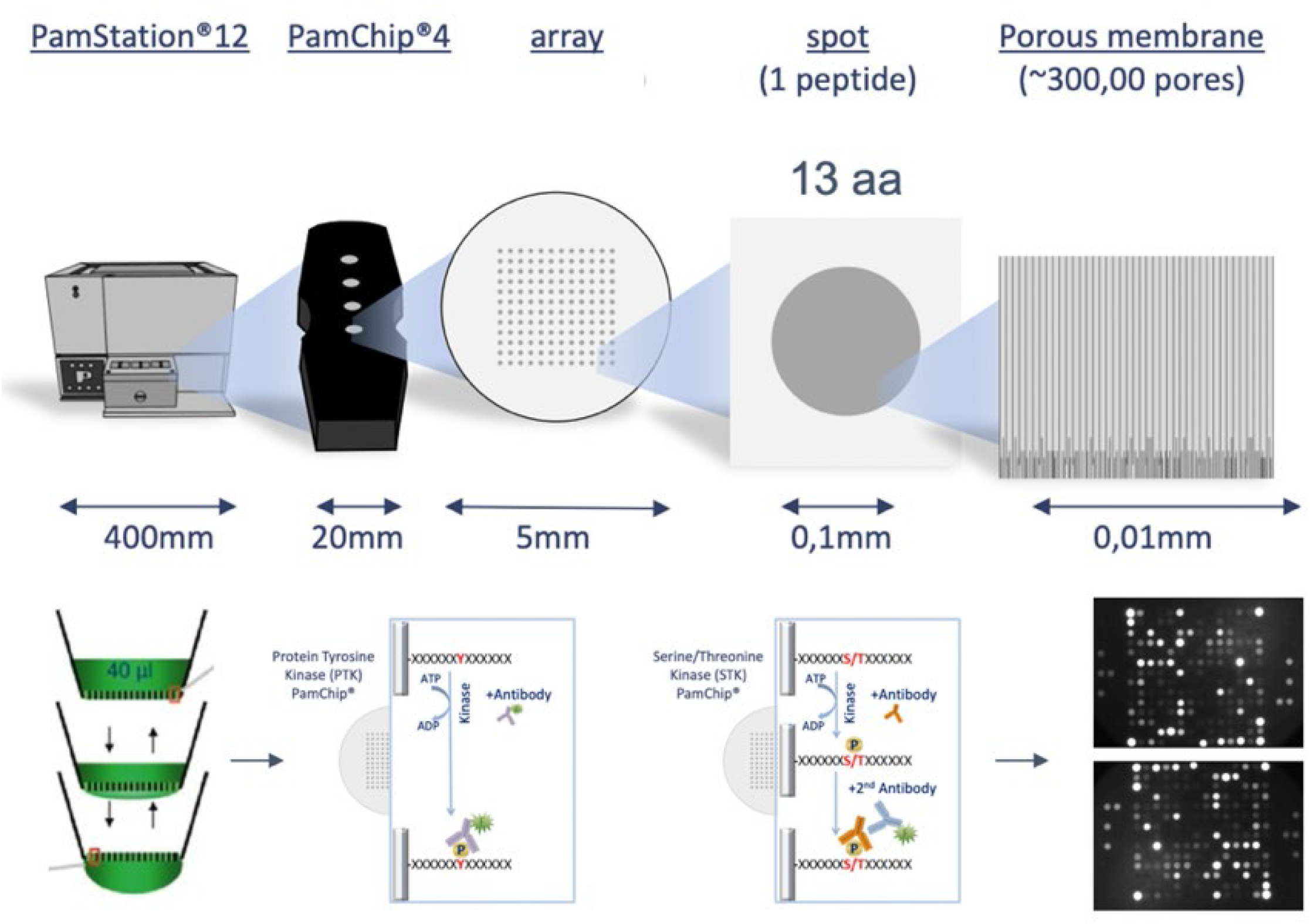
PamGene Platform Workflow. The arrays are spotted with reporter peptides, including controls, coupled to an activated aluminum oxide surface to create a 3-D structure facilitating interactions. During an experiment, the array is incubated with lysates of cells or tissue. The active kinases in the sample will phosphorylate their target on the array. Generic fluorescent labeled antibodies that recognize phosphorylated residues are used to visualize the phosphorylation in real time. This figure was reprinted with permission from PamGene International B.V..

**Supplemental Fig. 5.**
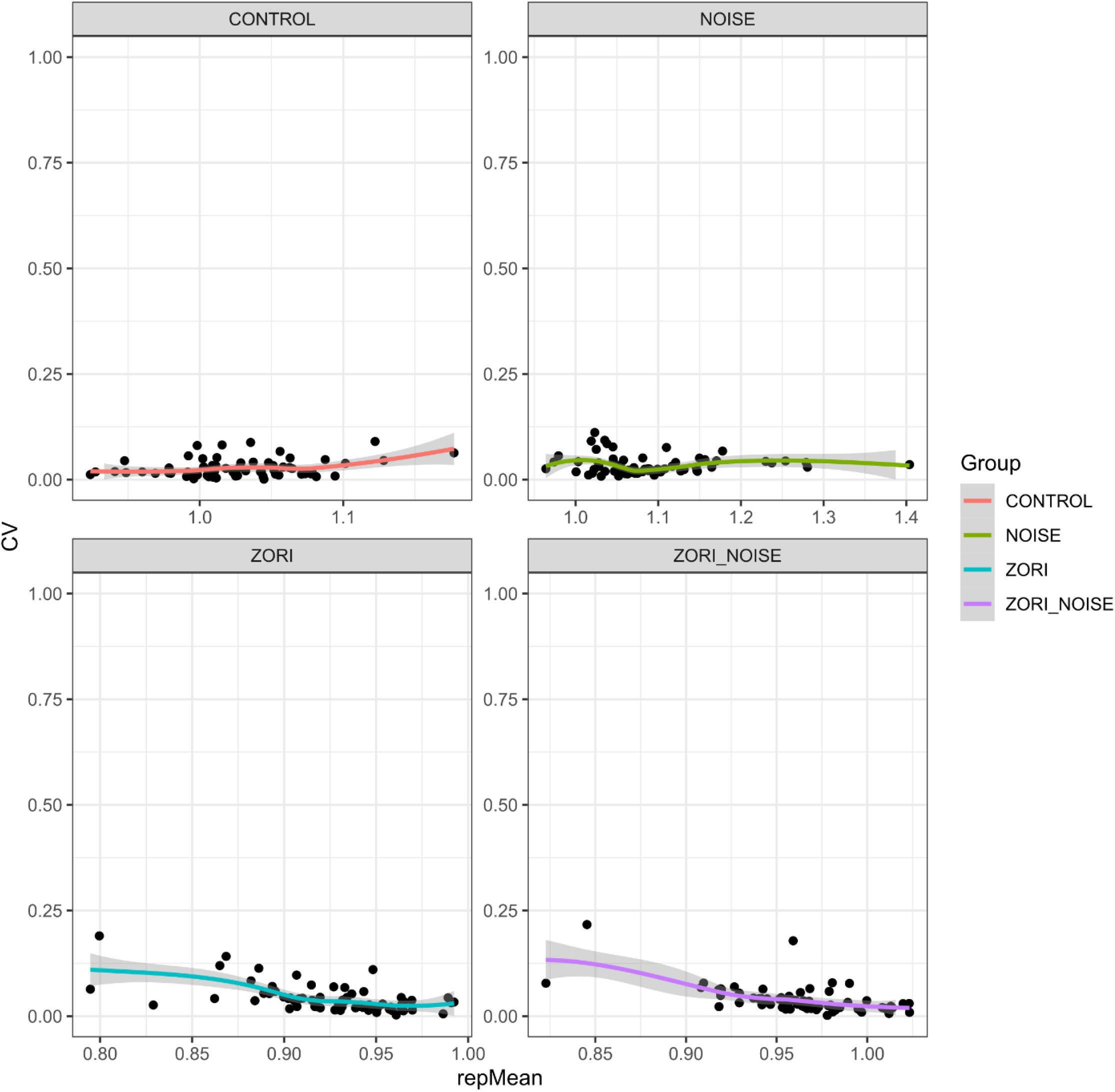
Coefficient of variation (CVs) of normalized signal intensities for each reporter peptide across control, zorifertinib (Zori), noise, and zorifertinib plus noise groups. Data show low variability across dozens of reporter peptides.

**Supplemental Fig. 6.**
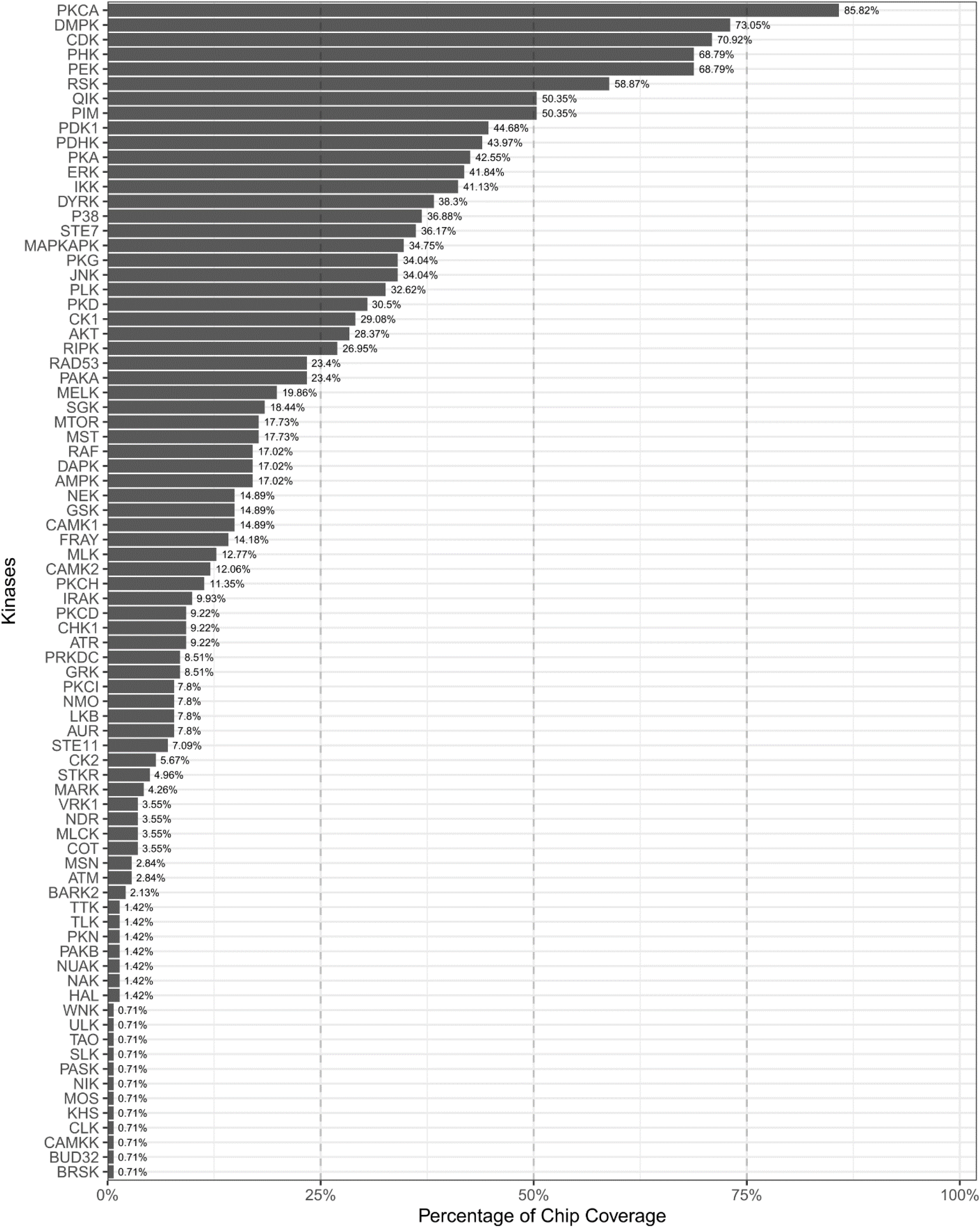
Coverage of protein kinase families based on assignments of protein kinases to specific STK chip reporter peptides generated by the kinome resampling analyses (KRSA) package. X-axis indicates percentage of peptides on chip that “map” to a kinase in a specific kinase family (Y-axis). There are 144 distinct reporter peptides on the STK chip.

**Supplemental Fig. 7.**
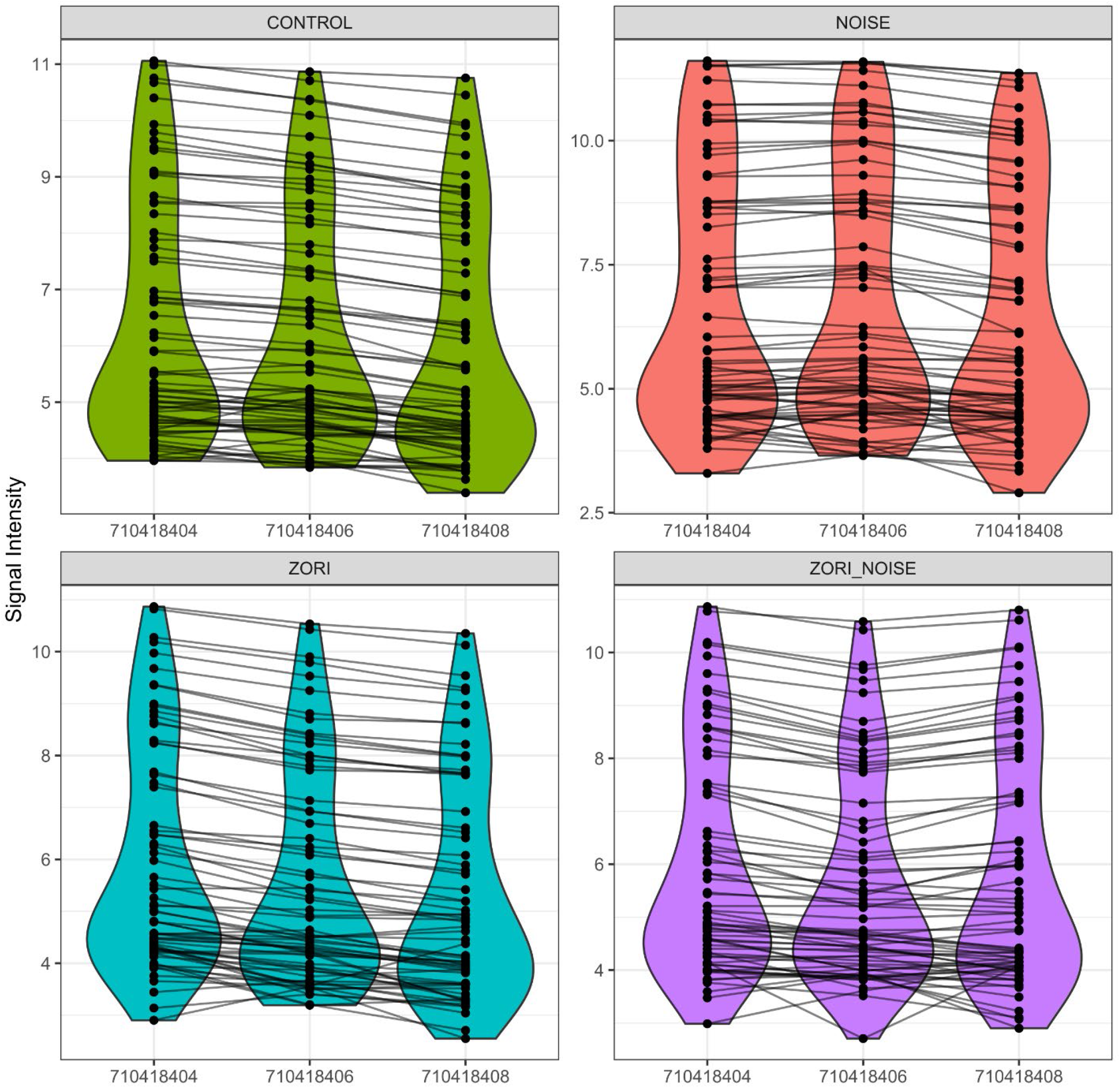
Distribution of normalized signal intensities of reporter peptides for control, zorifertinib (Zori), noise, and zorifertinib plus noise groups across three STK chips. Each group was included on each of three chips, providing n = 3 technical replicates. Each chip has 4 wells.

**Supplemental Fig. 8.**
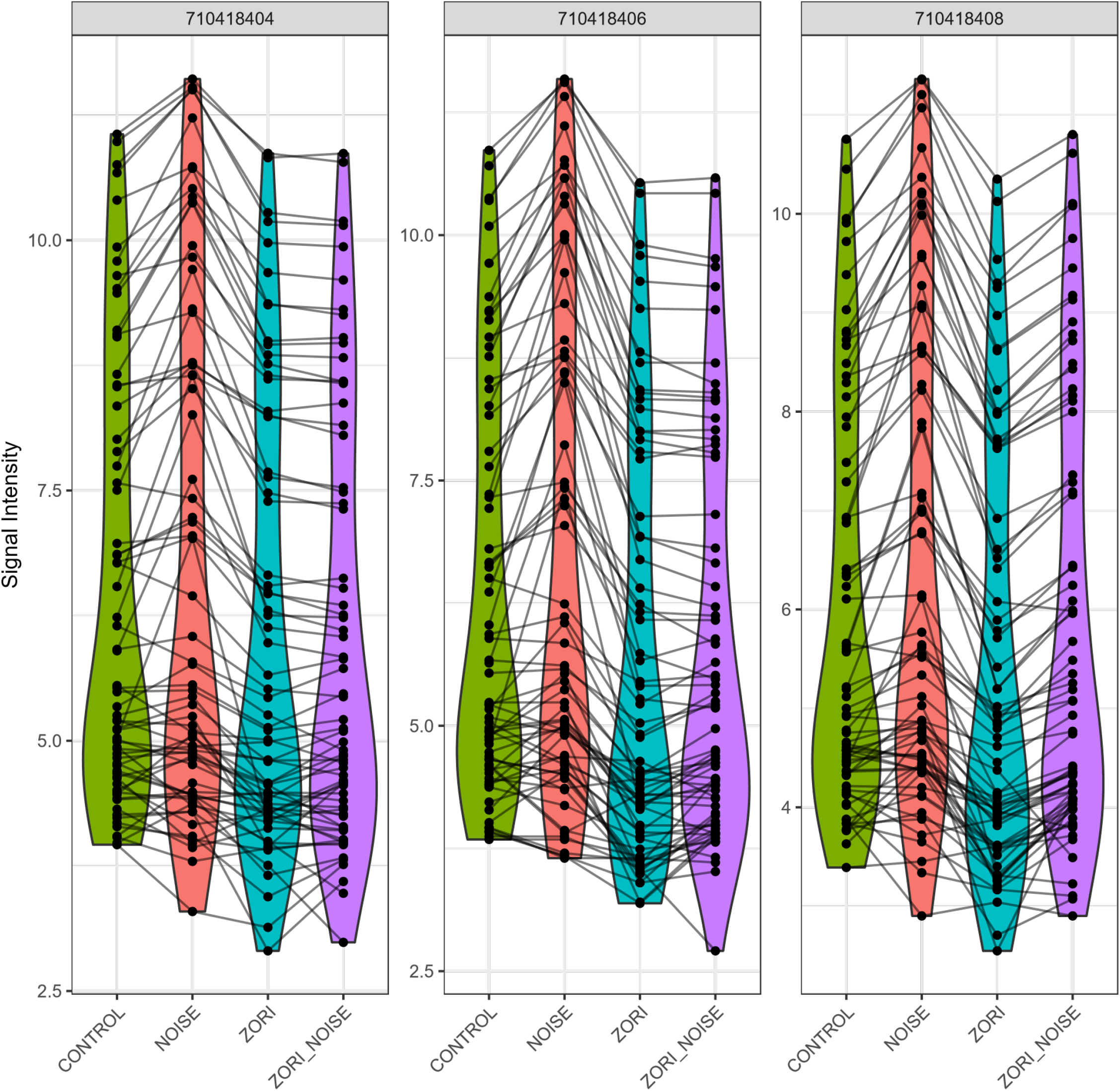
Distribution of normalized signal intensities of reporter peptides for control, zorifertinib (Zori), noise, and zorifertinib plus noise groups between three STK chips. Each group was included on each of three chips, providing n = 3 technical replicates. Each chip has 4 wells.

**Supplemental Fig. 9.**
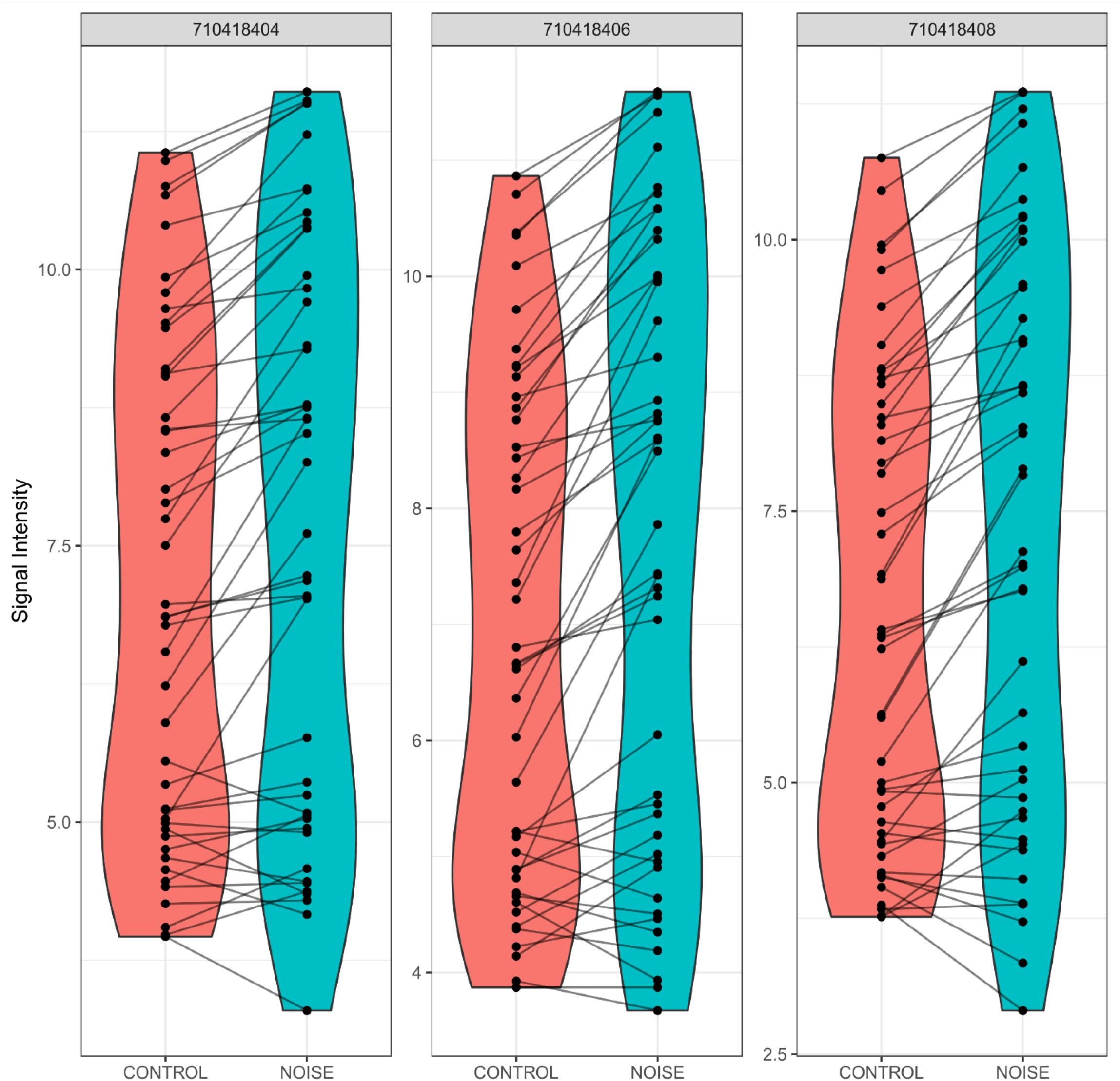
Distribution of normalized signal intensities of reporter peptides for noise versus control groups only across three STK chips.

**Supplemental Fig. 10.**
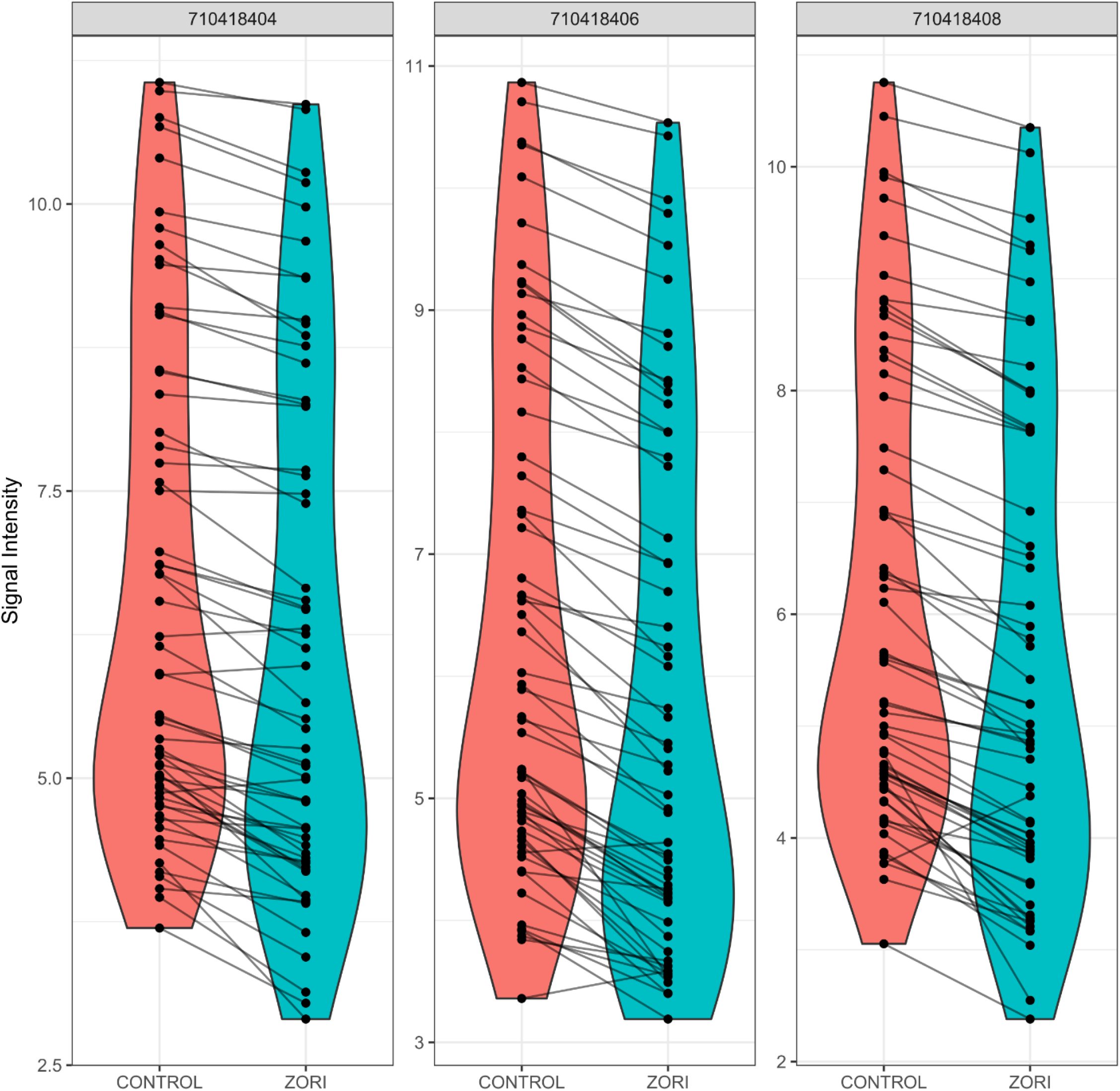
Distribution of normalized signal intensities of reporter peptides for control versus zorifertinib (Zori) groups only across three STK chips.

**Supplemental Fig. 11.**
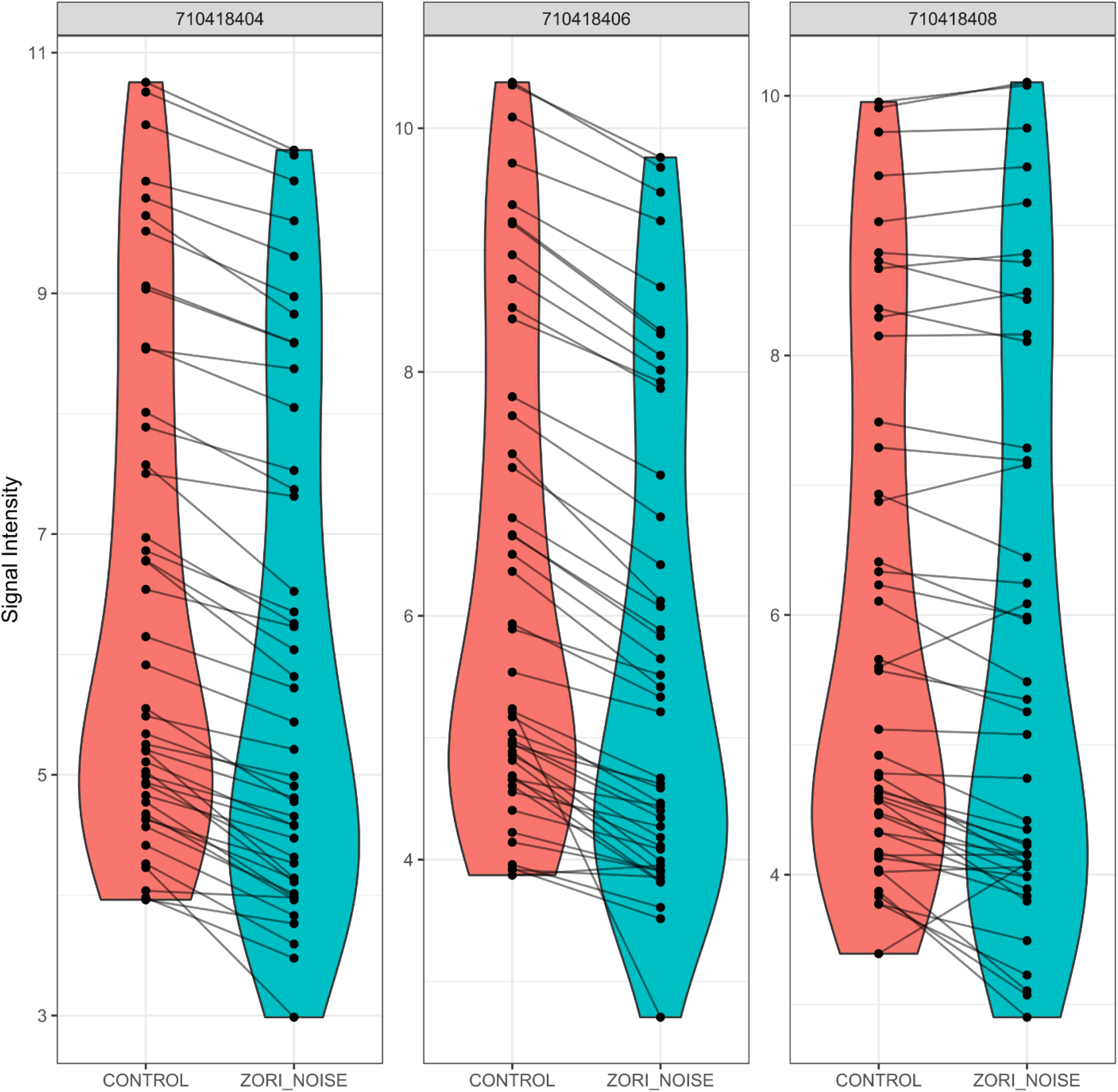
Distribution of normalized signal intensities of reporter peptides for control versus zorifertinib (Zori) plus noise groups only across three STK chips.

**Supplemental Fig. 12.**
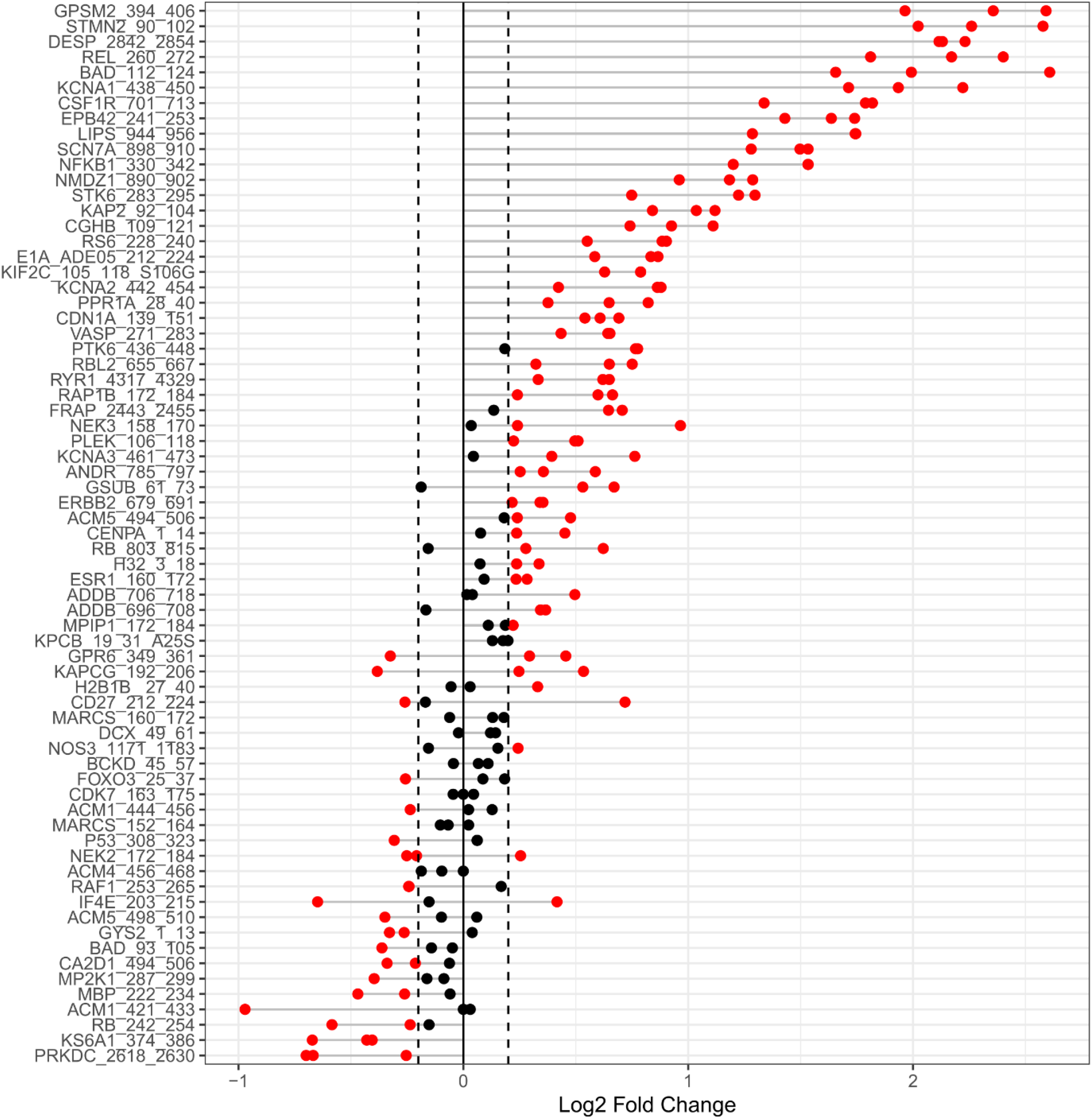
Waterfall plot of Log2 fold change (X-axis) in phosphorylation of reporter peptides (Y-axis) in control vs noise cochlear homogenate from three STL chips. Each dot represents the value for that peptide from one of the three chips for the same comparison. Red dots are outside the threshold (+/-0.15) considered meaningful for changes in kinase activity.

**Supplemental Fig. 13.**
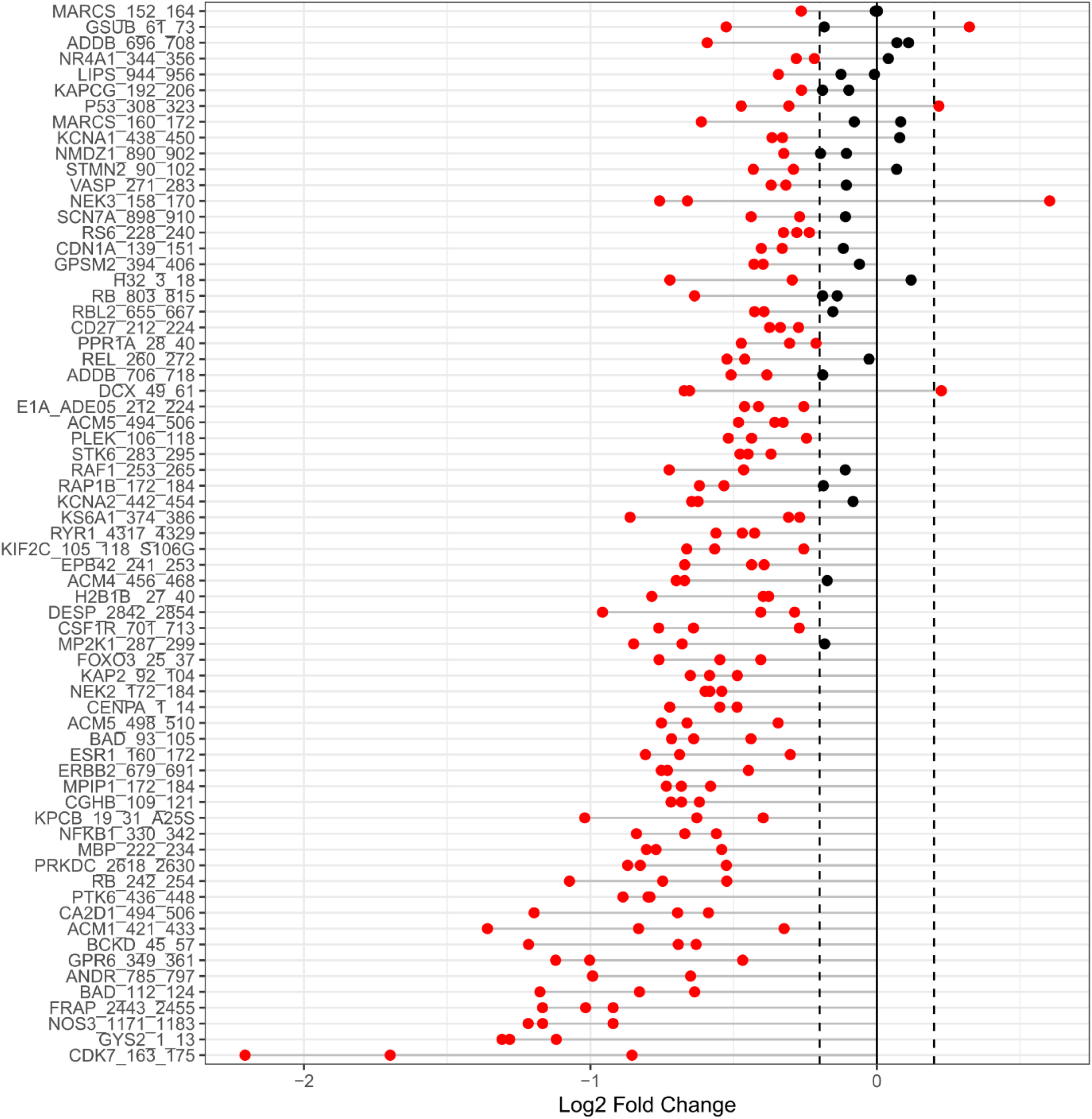
Waterfall plot of Log2 fold change (X-axis) in phosphorylation of reporter peptides (Y-axis) in control vs zorifertinib (Zori) cochlear homogenate from three STL chips. Each dot represents the value for that peptide from one of the three chips for the same comparison. Red dots are outside the threshold (+/-0.15) considered meaningful for changes in kinase activity.

**Supplemental Fig. 14.**
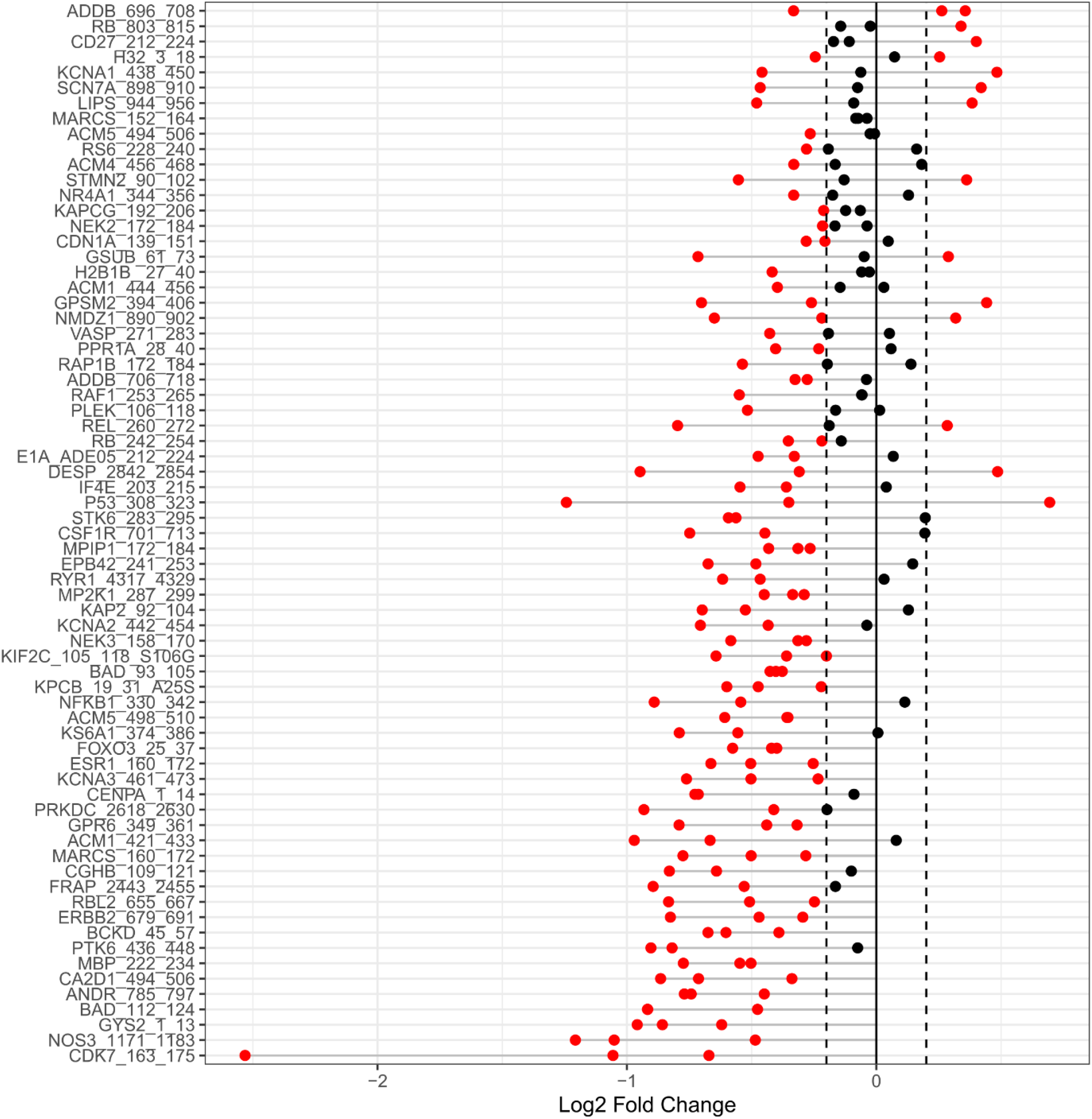
Waterfall plot of Log2 fold change (X-axis) in phosphorylation of reporter peptides (Y-axis) in control vs zorifertinib (Zori) plus noise cochlear homogenate from three STL chips. Each dot represents the value for that peptide from one of the three chips for the same comparison. Red dots are outside the threshold (+/-0.15) considered meaningful for changes in kinase activity.

**Supplemental Fig. 15.**
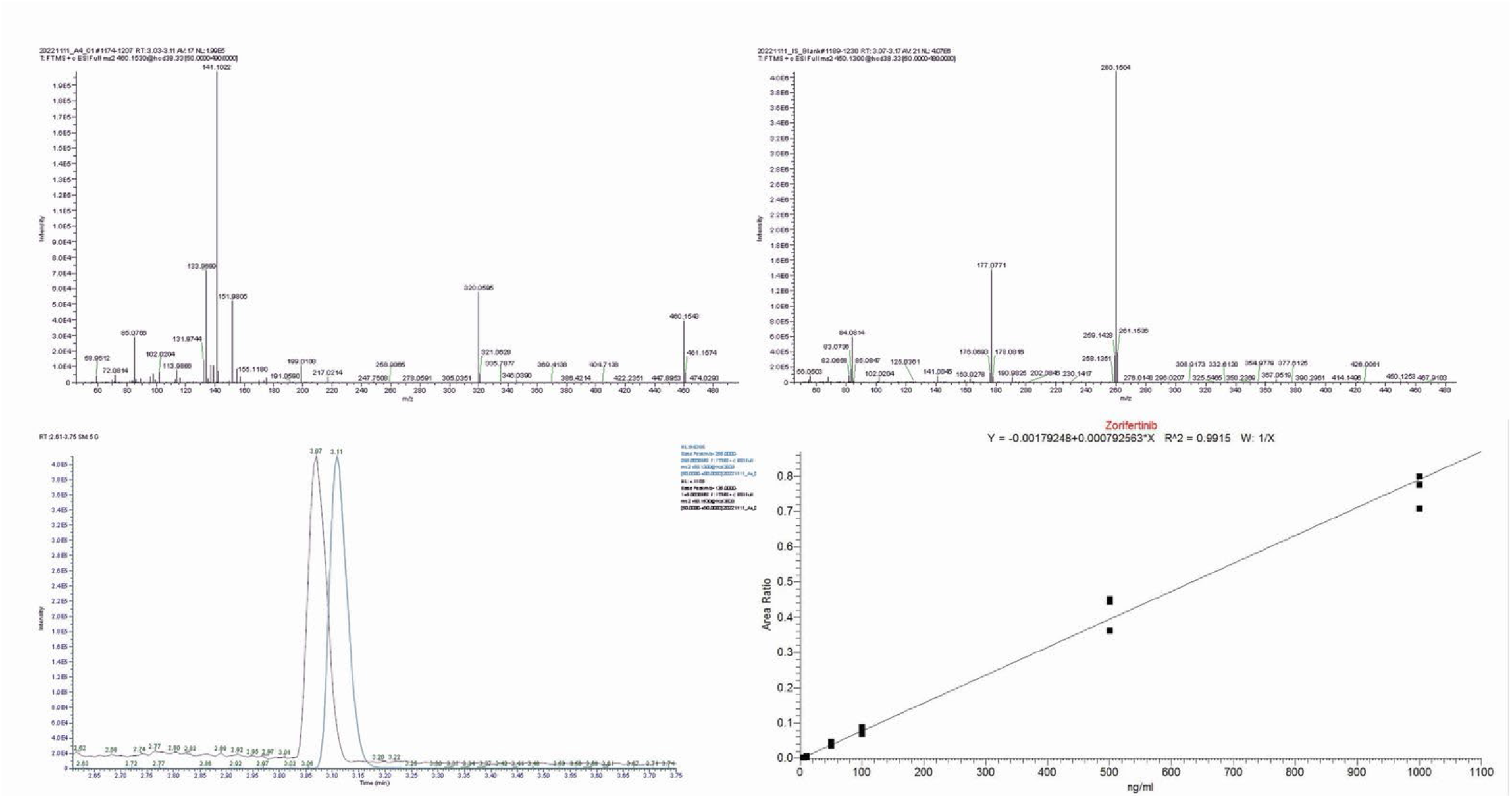
(A-B) MS2 spectra of Zorifertinib (A) and IS Crizotinib (B). (C) Sample chromatogram showing peaks for both zorifertinib (right) and IS (left). (D) Calibration curve linear regression.

